# Quantifying structural variants in chromosomes using landmark-based disparity

**DOI:** 10.1101/2024.10.30.620815

**Authors:** Ashwini V. Mohan, Anjali Goswami, Jeffrey W. Streicher

## Abstract

Chromosomal architecture has played a key role in the evolution of biodiversity. Detecting structural variants (SVs) on chromosomes has informed the study of speciation, sex determination, adaptation, and some of the earliest divergences in the tree of life. Here we present a computationally non-intensive approach, based on geometric morphometrics, that uses conserved DNA sequences as landmarks to quantify structural disparities of focal chromosomes across multiple species, individuals, or cell types. Based on two approaches, we show that this ‘geno-metric’ method can be applied at micro– and macroevolutionary scales to discover and diagnose SVs. Using human X-linked genes and ultraconserved elements as landmarks, we provide empirical demonstrations with amniote sex chromosomes, the *Drosophila virilis* group, and placental mammal genomes. Landmark-based structural disparity analysis effectively identifies chromosomal rearrangements and has parallels with traditional morphometrics regarding chromosome size, landmark orientation and landmark availability. Using simulations, we show that structural disparity inferred from ultraconserved elements is correlated with overall levels of chromosome evolution; an attribute which is consistent with observed disparity between and within mammalian orders. We found that the disparity patterns of SVs have significant phylogenetic signal, giving them broad importance for studying evolutionary biology. Structural disparity analyses are a valuable addition to the comparative genomic toolkit in that they offer an intuitive, rapid mechanism for detecting SVs associated with single copy genetic landmarks and the potential to reveal broader patterns of chromosome evolution related to expansions, contractions, rearrangements and phylogeny.

## Introduction

We are now firmly in the age of genomics with fully sequenced genomes commonplace for most organismal groups. Because of this, multi-species genomic comparisons are increasingly used in evolutionary and ecological research. This is even the case for organisms with relatively large nuclear genomes [1–3]. Excitingly, the comparison of whole genome sequences has offered valuable evolutionary insights into a range of topics including gene orthology [4], transposable element evolution [5], and adaptation to novel environments [6].

One important area of comparative genomics research is the study of chromosome architecture [7]. Detecting structural variants (SVs) of chromosomes is highly relevant to the evolution of biodiversity and disease [8]. From supergenes [9] to sex chromosomes [10], evolutionary shifts in chromosome structure are known to have a substantial influence on organismal phenotype. Recent research utilizing forward in time simulations has also indicated the important role of SVs in sustaining genomic adaptation [11]. Traditional methods for detecting SVs include chromosome banding and fluorescent in situ hybridization [12]. Several modern methods using genomic sequencing have also been developed including aberrant read alignment patterns, split reads, coverage changes, alignment of long-read sequences, and strand-seq [13–14]. Amongst sequencing-based methods, synteny assessment [15] is increasingly used to compare chromosome structure across multiple species with visual interpretation achieved through complex network diagrams, e.g. [16–19]. Most of these high-throughput DNA sequencing approaches for detecting SVs depend on comparisons to a single reference sequence and/or whole genome alignment, which can be subject to reference-bias and time consuming, respectively [20]. To complement existing methods, here we present a method for detecting SVs across chromosomes from multiple species or individuals simultaneously that is reference-free, computationally non-intensive, and does not require whole genome alignment. Based on the principles of morphometrics, our method is applied to chromosomes from high-quality genome assemblies and facilitates the study of SVs in ordinated micro– and macroevolutionary contexts.

Within the study of comparative morphology, structural variation in homologous structures across species is commonly termed ‘disparity’ [21–22]. Measures of morphological disparity are routinely used to explore the evolutionary dynamics of complex phenotypes, e.g. [23–24]. Structural disparity analysis of chromosomes is a disparity-based method that uses conserved landmark sequences to capture genomic variation across different species in a single analytical framework. Our ‘geno-metric’ method is based on a simple premise: chromosome structure is morphology (i.e., a shape, form, or structure) and there are well-validated methods for capturing morphological disparity across multiple species [23–25]. Furthermore, genomes are filled with homologous sequences (=landmarks) making them amenable for such analyses (e.g. [26]). To develop and scrutinize landmark-based structural disparity analyses, we analyzed three empirical datasets. First, we quantified SVs associated with sex chromosome evolution across amniotes using human X-linked BUSCO (Benchmarking Universal Single-Copy Orthologs) genes as landmarks which represents a scenario when the structure of a particular genomic/chromosomal feature is of interest [sensu 18). Second, we compared SVs related to a speciation-associated chromosome in Drosophila [27] using ultraconserved elements (UCEs; [28]) as landmarks which represents a scenario when chromosome-wide structure may be of interest. Third, we used genome sequences from placental mammals to develop a pipeline for discovering chromosomes that share homologous landmarks; a necessary step when chromosome homology is unclear or partial. We also used the placental mammal dataset to simulate SVs of chromosomes and determine the effects of indels and inversions on disparity measures. Finally, we tested all datasets for phylogenetic signals associated with disparity patterns of SVs.

## Results

Pipeline development. We developed two approaches for conducting structural disparity analysis, [1] using generic feature format (gff3) annotation files as input and [2] a de novo method that allows users to landmark and analyze whole chromosome sequences from genome assemblies (Fig. 1). The gff3-formatted pipeline allows users to extract genes from the chromosome of a single species in the gff3 file and then identify orthologs in the other species contained within the file. This approach is useful when chromosome homology is known, and focal landmarks are largely confined to single chromosomes. The de novo method begins by acquiring chromosome sequences from NCBI and then reference mapping landmarks to the assemblies using bwa [29]. We tested other approaches for reference mapping including minimap2 using default settings [30] but found bwa resulted in the most mapped landmarks per chromosome (Fig. S1). After orthologous landmarks are mapped, landmarked chromosomes are either prepared for downstream analysis (when chromosomal homology is known), or clustered based on the presence/absence of landmarks via metric multi-dimensional scaling (when chromosomal homology is not known). The latter option allows for the identification of chromosomes that share large numbers of landmarks, and thus likely large homologous genomic blocks. Following landmarking, both approaches have several steps in common including; removal of landmarks that are multi-copy or not shared across homologs, complementarity checks (5’-3’ versus 3’-5’), and exportation of landmark position information into a spreadsheet. The spreadsheet has rows representing species or individuals (one for each species or individual included in the input assemblies) and columns are landmark positions on a homologous chromosome set. The spreadsheet is then used as input for the disparity inference stage. Depending on user preference, absolute landmark positions can be used, or positions can be transformed using landmark bounding and/or chromosome size correction. Unless otherwise indicated, all steps in the pipeline occur using base functions in the R statistical package [31]. As with other disparity analyses, we use ordination techniques to reduce the dimensionality of variation in landmark position wherein the spreadsheet of landmark positions is analyzed using principal component analysis (Fig. 1).

**Figure 1.**
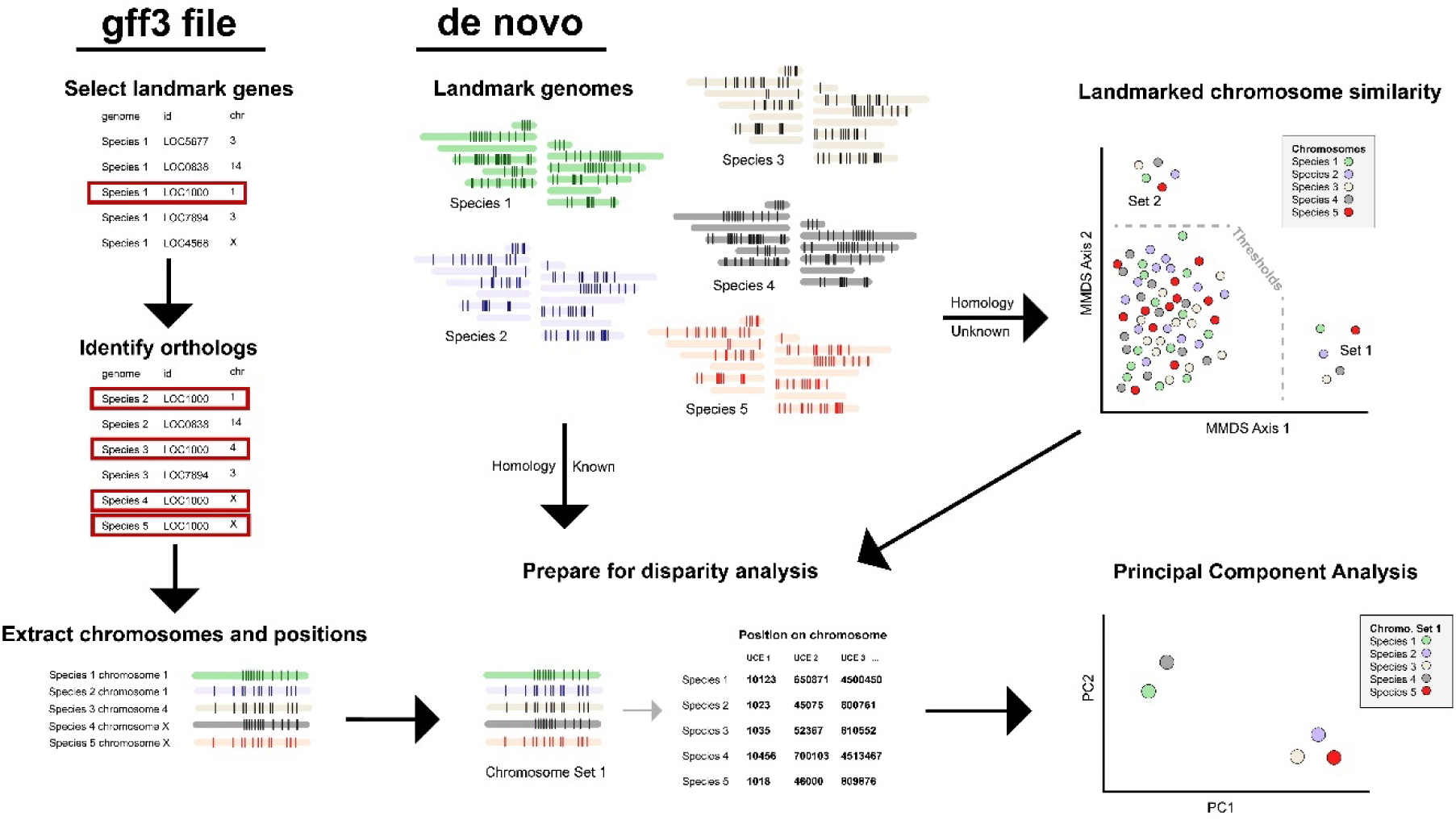
Schematic illustration of the landmarking pipeline for structural disparity analysis of chromosomes. When using generic feature format (gff3) annotation files, landmark genes are selected from a reference followed by identification of orthologs from other species in the file and extraction of chromosome information and landmark positions. In the de novo approach, the first step is reference mapping landmarks to chromosome-level genome assemblies of different species, individuals, or cell types. If the homology of chromosomes is known, then the chromosomes are further prepared for disparity analysis. If homology is unknown, a presence-absence matrix is used to identify chromosomes with similar landmark compositions using a gap-based threshold for similarity. Chromosome sets are prepared for disparity analysis by extracting chromosome position information for each landmark. The resulting data matrix is used for multivariate Principal Component Analysis to visualize disparity across species.

Structural disparity of human X-linked genes. Lovell et al. [18] examined 300 million years of vertebrate sex chromosome evolution using synteny-anchoring. We took their gff3-formatted annotation file and identified 53 human X-linked genes that could be used as landmarks on a set of single chromosomes for the focal taxa. We found that principal components (PCs) 1 and 2 explained 69.3% and 14.2% of structural variance, respectively. Using these first two axes, we found that major phylogenetic groups occupied distinct areas of geno-metric space (Fig. S2A). PC1 was significantly correlated with chromosome size (non-parametric Spearman’s test, rho = 0.70, p = 0.003), but not PC2 (rho = –0.27, p = 0.3043). Mean structural disparity was highest amongst those groups where the landmarks were found on sex chromosomes (placental mammals, marsupial mammals and squamate reptiles) compared to those where these markers are located on autosomal chromosomes (birds, monotreme mammals). When the data were analyzed using the reverse complement orientation of the chromosomes, we found that some chromosomes with clustered landmarks resulted in exaggerated PC scores for some taxa (e.g., chicken, *Gallus gallus*, Fig. S2B). We found that landmark bounding corrected for this methodological artifact regardless of the 5’ to 3’ orientation of the chromosomes (Fig. 2A, S2C, S2D). A size-corrected version of the analysis changed the patterns of disparity with birds being the most disparate group, followed closely by placental mammals (Fig. S2E). The size-corrected analysis was more robust to chromosome orientation with comparable results (Fig. S2F). Size-correction of the bounded version of the dataset generated indistinguishable results in the two directions of analysis (Fig. S2G, S2H). For comparison we generated a GENESPACE-style synteny link plot (Fig. 2B).

**Figure 2.**
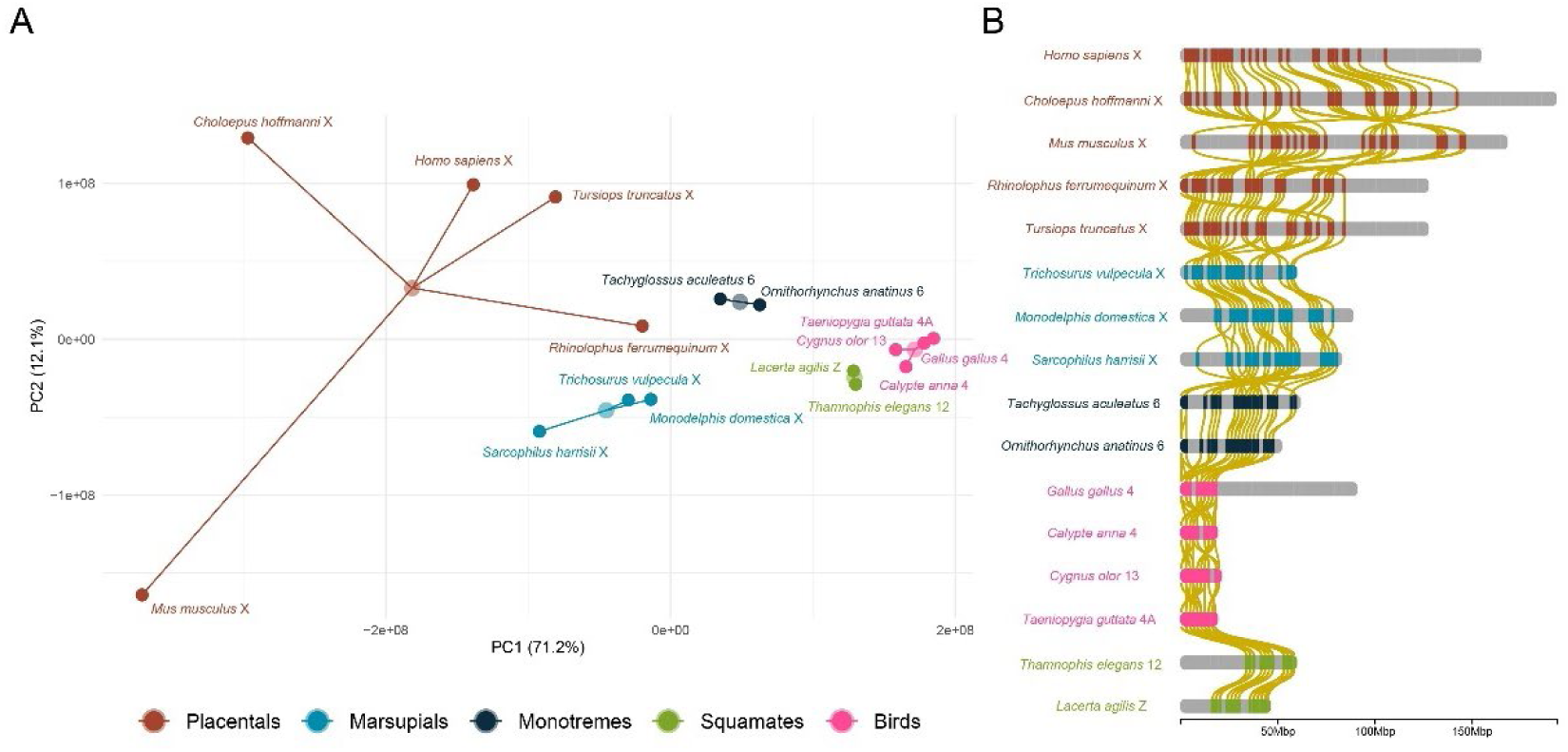
Structural disparity analysis of 53 human X-linked BUSCO gene landmarks in select amniote vertebrates studied by Lovell et al. (2022). Disparity analysis using bounded, absolute landmark positions reveals larger disparity in those groups where landmarks are found on the X chromosomes (placental mammals and marsupial mammals) than in those groups where landmarks are found on autosomes or Z chromosomes (A). This disparity is largely explained by chromosome size in that some of these groups maintain high synteny despite also having high structural disparity (B). Size-corrected disparity analyses capture different aspects of variation in landmark placement revealing birds as especially disparate after chromosome size is accounted for (Fig. S2E, F, G, H).

Structural disparity among closely related species of *Drosophila*. Poikela et al. [27] identified multiple chromosomal inversions that are associated with speciation patterns in the *Drosophila virilis* complex. We selected chromosome 5 for structural disparity analysis because there are known chromosomal inversions across the focal species [27]. We identified a set of 191 widely distributed dipteran UCEs [32] on chromosomes 5 for use as landmarks. Our structural disparity analysis revealed that three PC axes explained nearly 100% of the variance in landmark placement (PC1, 94%; PC2 5.2%; PC3, 0.07%). Patterns of disparity were highly polarized with PC1 separating *D. flavomontana* from the other species, PC2 separating *D. montana* from the other species, and PC3 scores separating *D. americana + D. novamexicana* from the two individuals of *D. virilis* (Fig. 3). Those individuals with highly conserved, syntenic UCE landmark placements occurred closest in geno-metric space. Notably, this was the case for the two individuals of *D. virilis* (Fig. 3A) and close relatives *D. americana* and *D. novamexicana* (Fig. 3B; [33]). In contrast, separation corresponded to distinct SVs. PC1 scores were largely explained by two blocks of UCE landmarks with substantially different placements in *D. flavomontana* compared to the other taxa (Fig. 3C). PC2 scores largely explained by a pericentric chromosomal inversion in *D. montana* relative to the other taxa (Fig. 3D). Importantly, when we compare the landmark orientations on *D. montana* and *D. flavomontana*, we see evidence (Fig. 3E) of the two blocks of UCE landmarks (PC1) and the pericentric inversion (PC2). PC3 scores were largely explained by a paracentric inversion that *D. virilis* has relative to both D. americana (Fig. 3F) and *D. novamexicana* (Fig. 3G). None of the three most explanatory axes were significantly correlated with chromosome size (PC1, rho = 0.029, p = 1.000; PC2, rho = –0.371, p = 0.4972; PC3, rho = 0.600, p = 0.2417).

**Figure 3.**
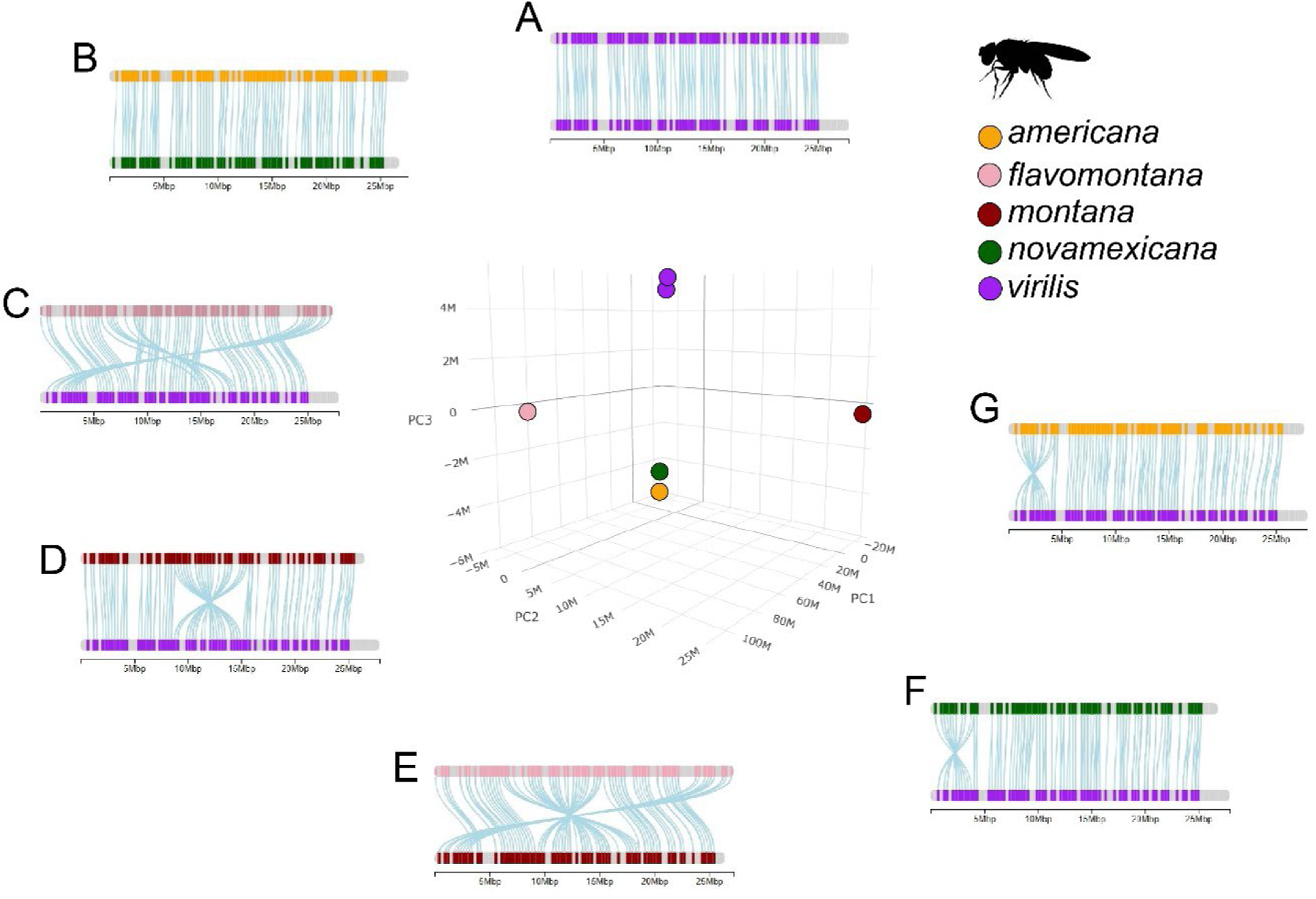
Structural disparity analysis using raw, absolute landmark positions of 191 ultraconserved elements (UCEs) on chromosome 5 of the *Drosophila virilis* group, different colors correspond to different species in the complex. Synteny link maps around the main plot demonstrate what separation in geno-metric space corresponds to, in terms of landmark placement on chromosome 5. When samples are close to one another in space they are highly syntenic (A, B). Different axes correspond to discrete differences in landmark placement including two translocations (C), a pericentric inversion (D, E) and paracentric inversion (E, F). An ‘M’ was arbitrarily added by plotly (91) to values on component axes because they are less than one. Synteny link maps were generated using chromoMap [35).

De novo homology discovery and structural disparity analysis using whole genome assemblies. Over long periods of evolutionary time, genome structure may change, substantially eroding the homology of chromosomes. Despite this, large genomic regions often remain evolutionarily intact on single chromosomes (e.g., [16]) among which structural disparity can be explored. We used the chromosome clustering module of our de novo pipeline to identify several chromosome sets containing homologous tetrapod UCEs [34] from 26 genome assemblies of five placental mammal orders (Artiodactyla, Carnivora, Perissodactyla, Primates, and Rodentia; Table 1). The results of the 10-axis chromosome clustering revealed that score thresholding could be used to identify eight sets of mutually exclusive chromosomes that (a) were highly similar in their UCE compositions and (b) contained only a single chromosome per species in the dataset (Fig. 4A, Table 2). Chromosome similarity and threshold gaps decayed to after the second axis with not all sets containing 26 species and some instances of overlap with chromosomes belonging to other sets (e.g. *Giraffa tippelskirchi* chromosomes in sets 7 and 8, Fig 4A, Table 2). Axes 9 and 10 did not reveal any novel chromosome clusters associated with UCE landmark content (Table 2). Viewing axes as bivariate plots made some clustering patterns clearer, especially in axis 4 (Fig. 4B, C). Given (a) the clear separation of chromosomes sets 1 and 2, and (b) that protein content was significantly more similar within these chromosome sets than between them (W = 6742, p < 0.0001, nonparametric Wilcoxon tests, Fig. S3) we used these chromosomes as input for a structural disparity analyses based on SVs of their UCE landmarks.

**Figure 4.**
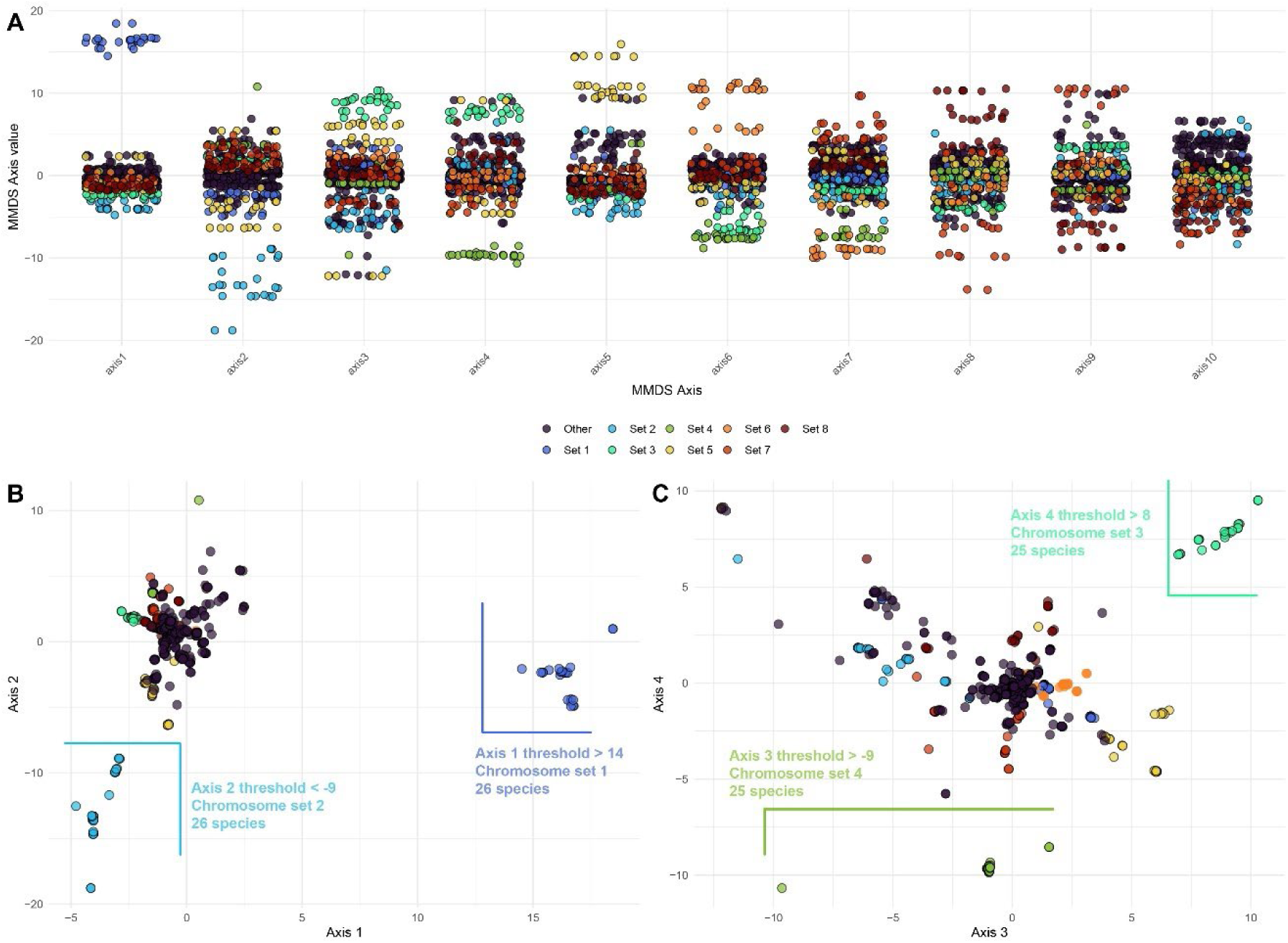
Metric multi-dimensional scaling (MMDS) of 567 landmarked placental mammal chromosomes based on patterns of ultraconserved element presence and absence. These chromosomes represent the genomes of 26 species (Table 1). MMDS axis thresholds are designated to establish ‘chromosome sets’ which contain a chromosome for most species included in the dataset. These chromosome sets likely contain mutually exclusive large homologous genomic blocks that are evolutionarily conserved and can be compared via structural disparity analysis. An initial assessment of 10 axes revealed eight sets of chromosomes that may be ideal for structural disparity analysis (A). Visualizing the clustering patterns in bivariate plots can assist with determining axis thresholds for defining the sets (B, C).

**Table 1.**
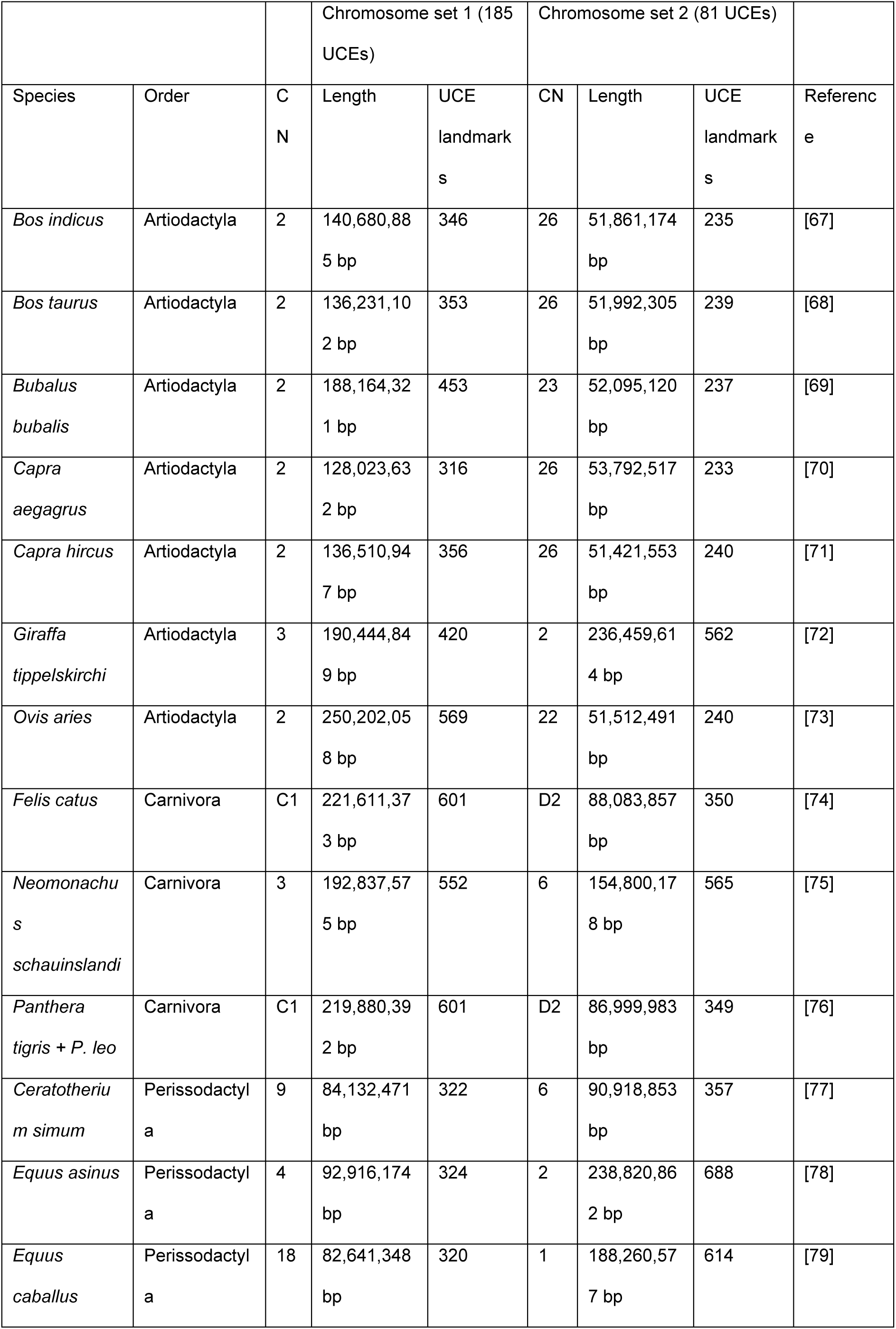
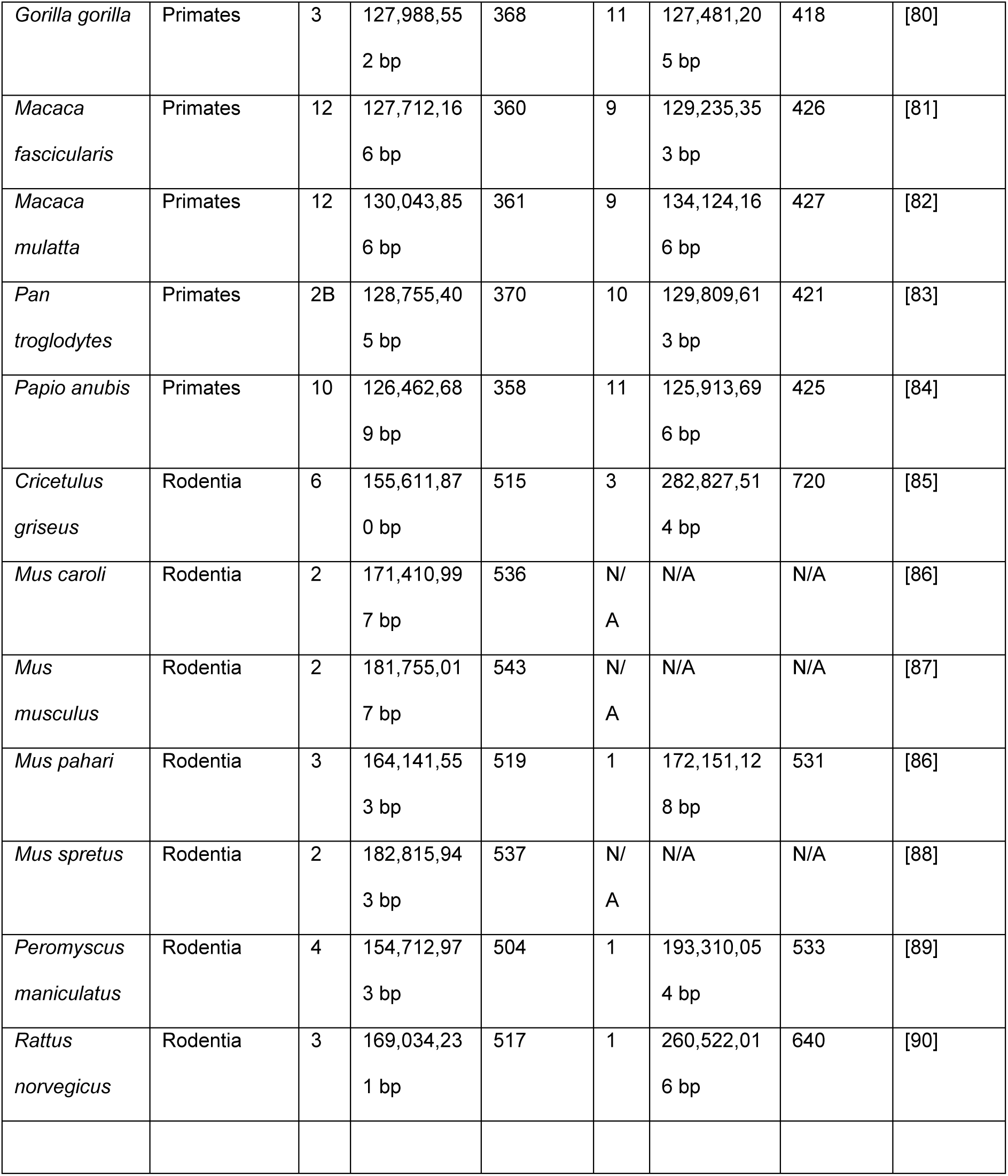
Chromosome sets identified from genome assemblies used to conduct landmark-based estimates of structural disparity. Those chromosomes appearing in bold text were used in the simulation/validation section of our study. CN = chromosome number (= reference number used in genomic assemblies), bp = base pairs, UCE = ultraconserved elements. Following removal of an outlier landmark, the chromosome sets had 185 and 81 landmarks, respectively.

**Table 2.** Axis thresholding based on metric multidimensional scaling (MMDS) of 567 placental mammal chromosomes from 26 species. Total number of species are unique chromosomes found to cluster on a given axis that are mutually exclusive when compared to clustering patterns of other axes. Threshold value is the smallest axis value identified in each group that is used to define the different clusters. Threshold gap value is the amount of space between the chromosome set and other chromosomes. An asterisk indicates two chromosomes sets where Macaca mulatta and/or *Giraffa tippelskirchi* chromosomes from other sets were within the threshold range. See Figure 4 for additional information.

| MMDS Axis | Number of species | Axis threshold | Threshold gap |
| --- | --- | --- | --- |
| 1 | 26 | 14.52 | 12.03 |
| 2 | 26 | -8.90 | 2.52 |
| 3 | 25 | 6.96 | 0.38 |
| 4 | 24 | -8.54 | 2.78 |
| 5 | 26 | 9.44 | 0.03 |
| 6 | 26 | 3.40 | 0.72 |
| 7 | 26* (*Giraffa* set 5, *Macaca* other) | 4.21 | 0.01 |
| 8 | 24* (*Giraffa* set 2) | 3.60 | < 0.01 |
| 9 | 0 | 0 | 0 |
| 10 | 0 | 0 | 0 |

Landmarking of UCES on the two chromosome sets resulted in 186 (set 1) and 82 (set 2) landmarks, respectively. Using absolute landmark positions (Figs. S4A, S5A), we found that most of the variation in both chromosome sets was explained by PC1. In chromosome set 1, PC1 described 98.9% of the variation and in chromosome set 2, PC1 described 99.4% of the variation. We discovered that in both chromosome sets, PC1 was significantly correlated with chromosome size (set 1, rho = –0.65, p < 0.0001; set 2, rho = 0.76, p < 0.0001) when using absolute size. However, this was not the case for PC2 (set 1, rho = 0.19, p = 0.3308; set 2, rho = 0.37, p = 0.066). Bounding the landmark positions increased the amount of variation explained by PC1 (chromosome set 1, 99.2%, Fig. S4C; chromosome set 2, 99.9%, Fig. S5C), and produced unexpected patterns with several notable outliers across the first two PCs (e.g. Cricetulus griseus, Fig. S5C). Size-corrected analyses of chromosome set 1 resulted in each mammal order occupying a unique region of geno-metric across PCs 1 and 2 (Fig. S4E), but this was not the case in chromosome set 2 where orders largely overlapped (Fig. S5E). Size-correction using bounded landmarks resulted in less variation explained by PC1 for both chromosome sets (chromosome et 1, 89.2%, Fig. S4G; chromosome set 2, 91.6%, Fig. S5G), and similar outlier patterns that were observed in the raw bounded position analysis. Mapping of component loadings onto chromosome plots revealed a single landmark (UCE 6948) that was heavily loaded on PC1 and PC2 of both chromosome sets (Fig.S6). Removing this outlier landmark changed the inferred structural disparity patterns across the PCAs with mammalian orders mostly occupying distinctive regions of geno-metric space in both chromosome set 1 (Fig. S4B, D, F, H) and, apart from a single size-corrected analysis (Fig. S5F), chromosome set 2 (Fig. S5B, D, H).

Following the removal of the outlier landmark, we found that PC1 explained most variation across the analyses. When using raw, absolute landmark positions PC1 explained 99.9% of the variance for both chromosome sets (Fig. S7A, S8A). Unlike the analyses including the outlier landmark, using bounded landmarks reduced the amount of variation explained on PC1 (95.3%, chromosome set 1, Fig. S7B; 98.6% chromosome set 2, Fig. S8B) compared to the raw positions. PC2 scores in the bounded analyses explained 2.0% (chromosome set 1) and 0.8% (chromosome set 2). Notably, the bounded landmark PC1 and PC2 separated all mammalian orders into distinct regions of geno-metric space. Size-correction via chromosome size resulted in different disparity patterns in both datasets with higher disparity within Artiodactyla in the chromosome set 1 analysis (Fig. S7E) and substantial overlap across orders in the chromosome set 2 analysis (Fig. S8E). The size-corrected, bounded analyses featured tighter clustering patterns within mammalian orders than the raw, size-corrected analyses in both chromosome sets (Figs. S7G, S8G). As observed in the X-linked BUSCO analyses, we found that using raw landmark positions on the reverse complemented version of the chromosome resulted in outlier taxa (*Macaca mulatta*, *Mus musculus*, chromosome set 1, Fig. S7B; *Equus asininus*, *E. caballus*, *Cricetulus griseus*, *Mus pahari*, *Peromyscus maniculatus*, *Rattus norvegicus*, chromosome set 2, Fig. S8B). However, unlike the X-linked BUSCO analyses, these outliers were also observed in the size-corrected analyses (Fig. S7F, S8F). Reverse complemented versions of the raw and size-corrected bounded analyses were more consistent with their equivalent original analyses (Fig. S7D, E, S8D, E). As observed in the X-linked BUSCO analyses, the amount of variance explained by PCs 1 and 2 in the size-corrected, bounded analyses was identical for both directions of analysis.

Based on our results, we selected the raw, bounded analysis with outlier landmark removed for further interpretation and analysis (Fig. 5). Interestingly, although they explained a small proportion of the overall structural variation, PC3 (0.5%, chromosome set 1, 0.5% chromosome set 2) and PC4 (0.1% chromosome set 1, 0.09% chromosome set 2) still clustered mammalian orders in largely distinct regions of geno-metric space for both chromosome sets (Fig. S9). However, *Capra aegragus* was an outlier on PC3 of chromosome set 1 (Fig.S9A). Synteny analysis of landmarks using chromoMap [35] to visualize structural differences captured by UCE landmarks revealed that certain chromosomal regions are more conserved as indicated by low PC loading values of landmarks into PC1 (blue links, Fig. 6), compared to other regions where landmarks show higher differences in their placement (maroon links, Fig. 6). Unlike the *Drosophila* example where landmark inversions explained most structural disparity (Fig. 3), insertions and deletion events between the UCE landmarks account for the structural disparity observed in the two placental mammal chromosomes sets (Fig. 6).

**Figure 5.**
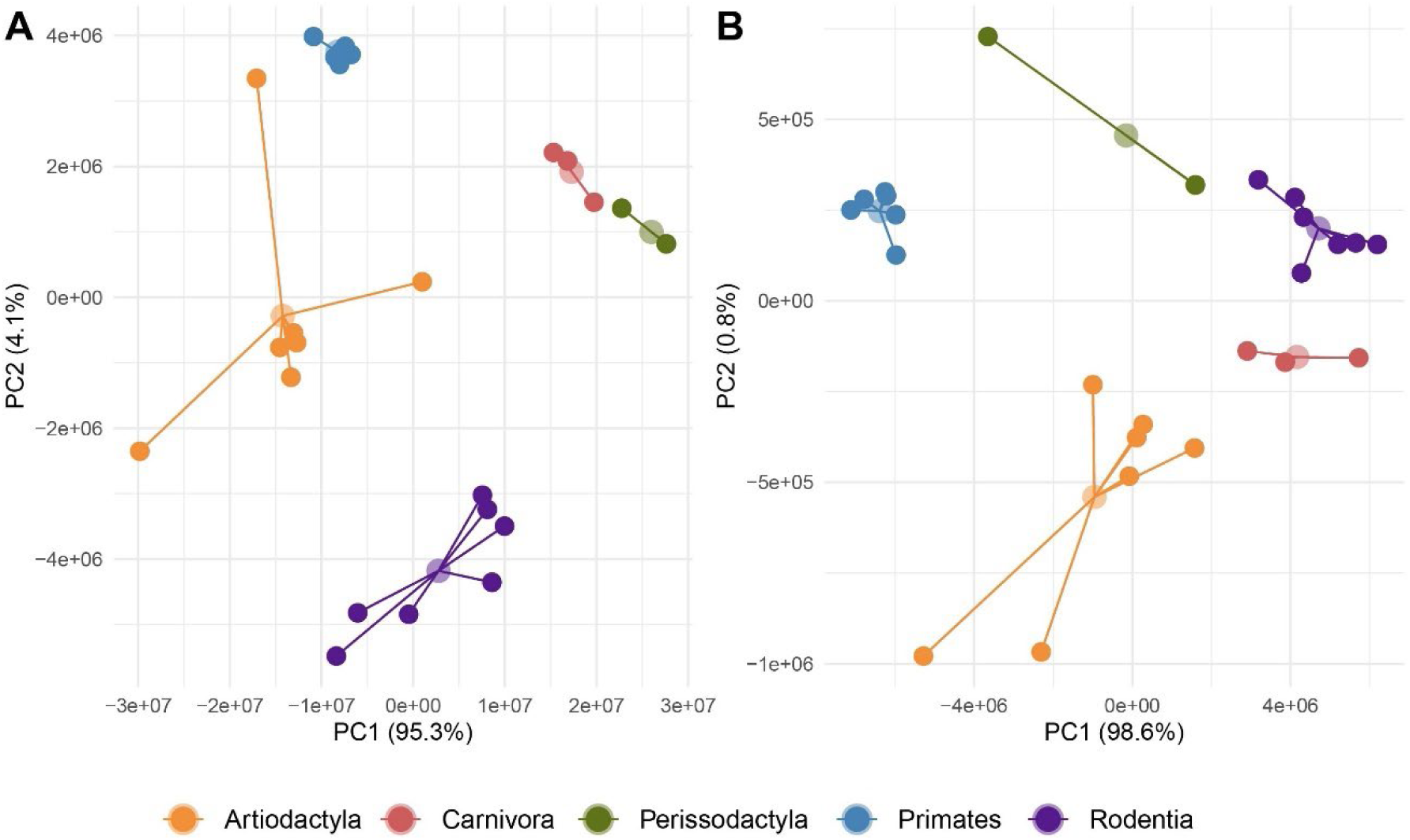
Ordination results from principal component (PC) analysis of UCE landmark positions in two sets of chromosomes from 26 species of placental mammals using bounded raw positions after the removal of an outlier landmark (see text). Representatives for five orders (Artiodactyla, Carnivora, Perissodactyla, Primates and Rodents) were included (Table1). The depicted structural variation is inferred from 185 (chromosome set 1, A) and 81 (chromosome set 2, B) landmarks, respectively. In both datasets, the first two PCs explained > 99% of the variation. Comparison of PCs 3 and 4 for both chromosome sets is provided in Figure S9.

**Figure 6.**
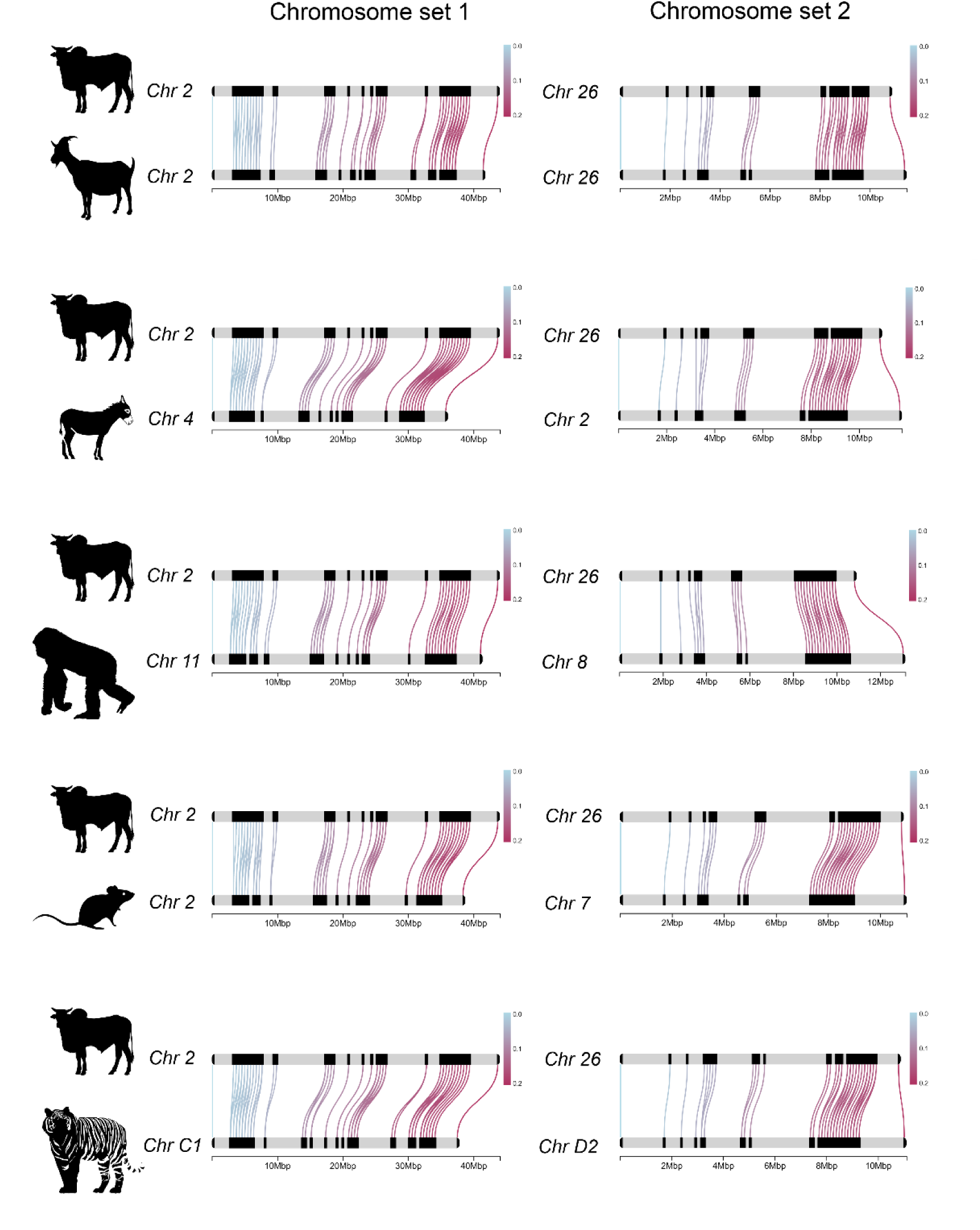
Select synteny link maps between *Bos indicus* and other placental mammals using bounded, absolute landmarking. Principal component loadings for various landmarks are indicated by the color of the links. The chromosome number (Chr) is indicated next to each species’ silhouette. The left and right columns are results from chromosome set 1 and set 2, respectively. Comparisons are (from top to bottom) between *Bos indicus* to *Capra aegagrus* (Artiodactyls), *Bos indicus* and *Equus asinus* (Perissodactyla), *Bos indicus* and *Gorilla gorilla* (Primates), *Bos indicus* and *Mus musculus* (Rodentia), and *Bos indicus* and the liger (*Panthera tigris + P. leo*, Carnivora). Synteny link maps were generated using chromoMap [35].

Structural disparity increases through time. In the simulation study, > 99% of the variation in landmark placement was explained by PC1 with PC2 explaining 0.2% of the variation. As predicted, pairwise increase in disparity (ID) increased with the amount of simulated evolution confirming that, at least under neutral (random) conditions, disparity measures should correspond to relative amounts of genomic evolution (Fig. 7A-D). We compared the variance of log-transformed ID scores for indel versus inversion simulations using two-way ANOVAs. We found that on both axes, inversions caused a significantly higher increase in disparity than indels (PC1, F =2.88, p = 0.0251; PC2, F = 3.34, p = 0.0121; Fig. S10). Using empirical data, we found that inter-order disparity was significantly higher than intra-order disparity for both PC1 (chromosome set 1, W = 63997, p < 0.0001; chromosome set 2, W = 67457, p < 0.0001, Fig. 7E, F) and PC2 (chromosome set 1, W = 67997, p < 0.0001; chromosome set 2, W = 61201, p < 0.0001, Fig. 7G, H). We found that protein-coding content tended to be more similar within mammalian orders than between them, however, this was not a significant result (W = 499.5, p = 0.0722; nonparametric Wilcoxon tests; SI Appendix, Fig. S11).

**Figure 7.**
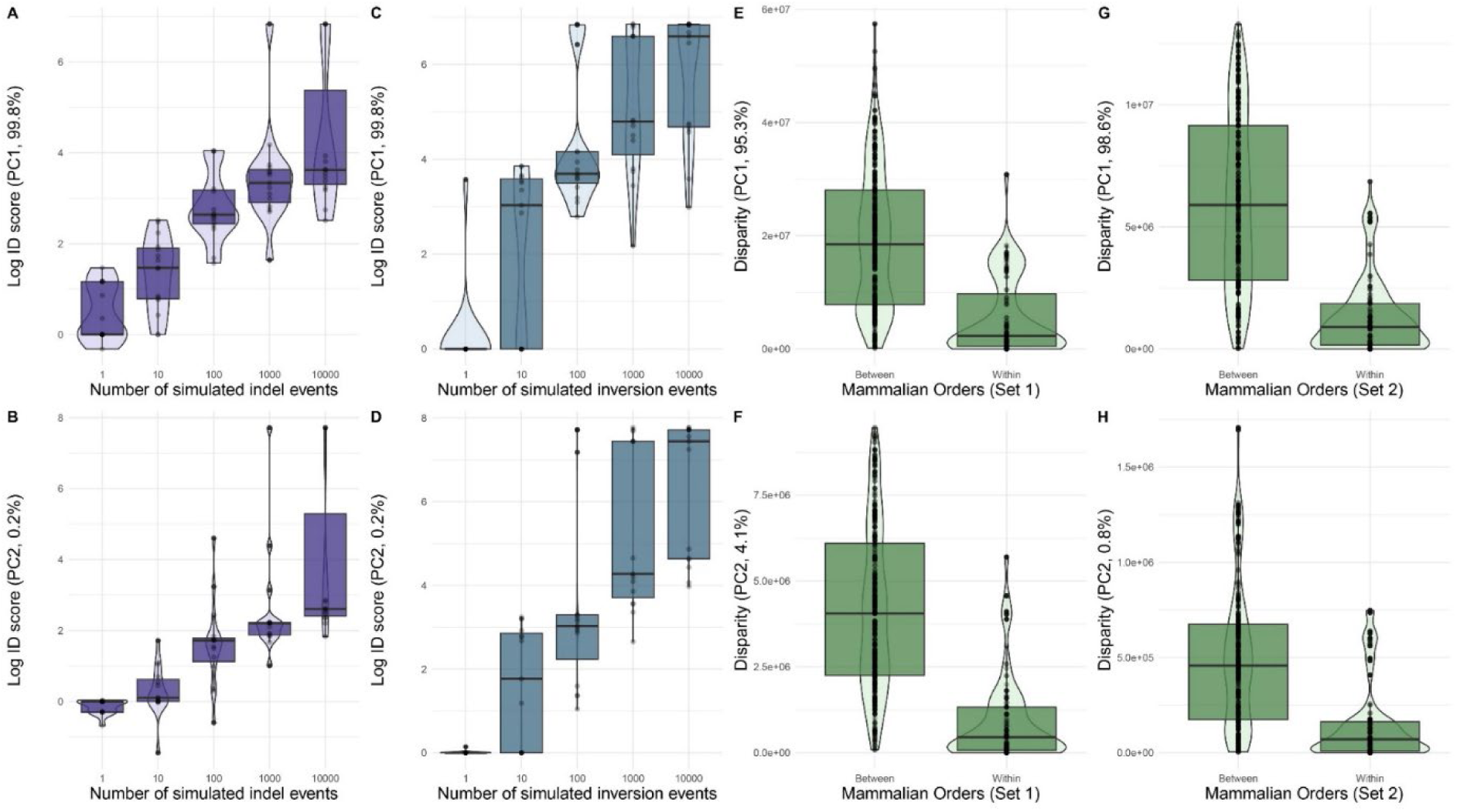
Log-transformed increase in disparity (ID) scores inferred from principal component (PC) analysis of ancestral and descendent insertion-deletion simulations for PC1. (A) and PC2 (B). Log-transformed ID scores inferred from inversion simulations for PC1 (C) and PC2 (D). Insertion-deletion simulation results are depicted in purple and inversion simulation results are depicted in blue. Observed empirical disparity scores for within and between mammalian orders for PC1 (E) and PC2 (F) of chromosome set 1 and PC1 (G) and PC2 (H) of chromosome set 2 are depicted in green.

Phylogenetic signal of structural variants. We found that the structural disparity captured by the most explanatory axes in all datasets had significant phylogenetic signal in 58% of the test cases (14/24 tests, Table 3). This included some cases where PC3 (describing < 1% of the overall variation) had significant phylogenetic signal (e.g. X-linked BUSCO genes, placental mammal UCE chromosome sets). Values of K and λ were also high (indicative of putative phylogenetic signal) in several non-significant tests. Ancestral state estimations of disparity revealed an increase in structural disparity related with the evolution of an X chromosome in the ancestor of placental and marsupial mammals (Fig. 8A), the evolution of a pericentric inversion on chromosome 5 in the *Drosophila virilis* group (Fig. 8B), and phylogenetically associated patterns of structural disparity increase or decrease in placental mammals (Fig. 8C, D).

**Figure 8.**
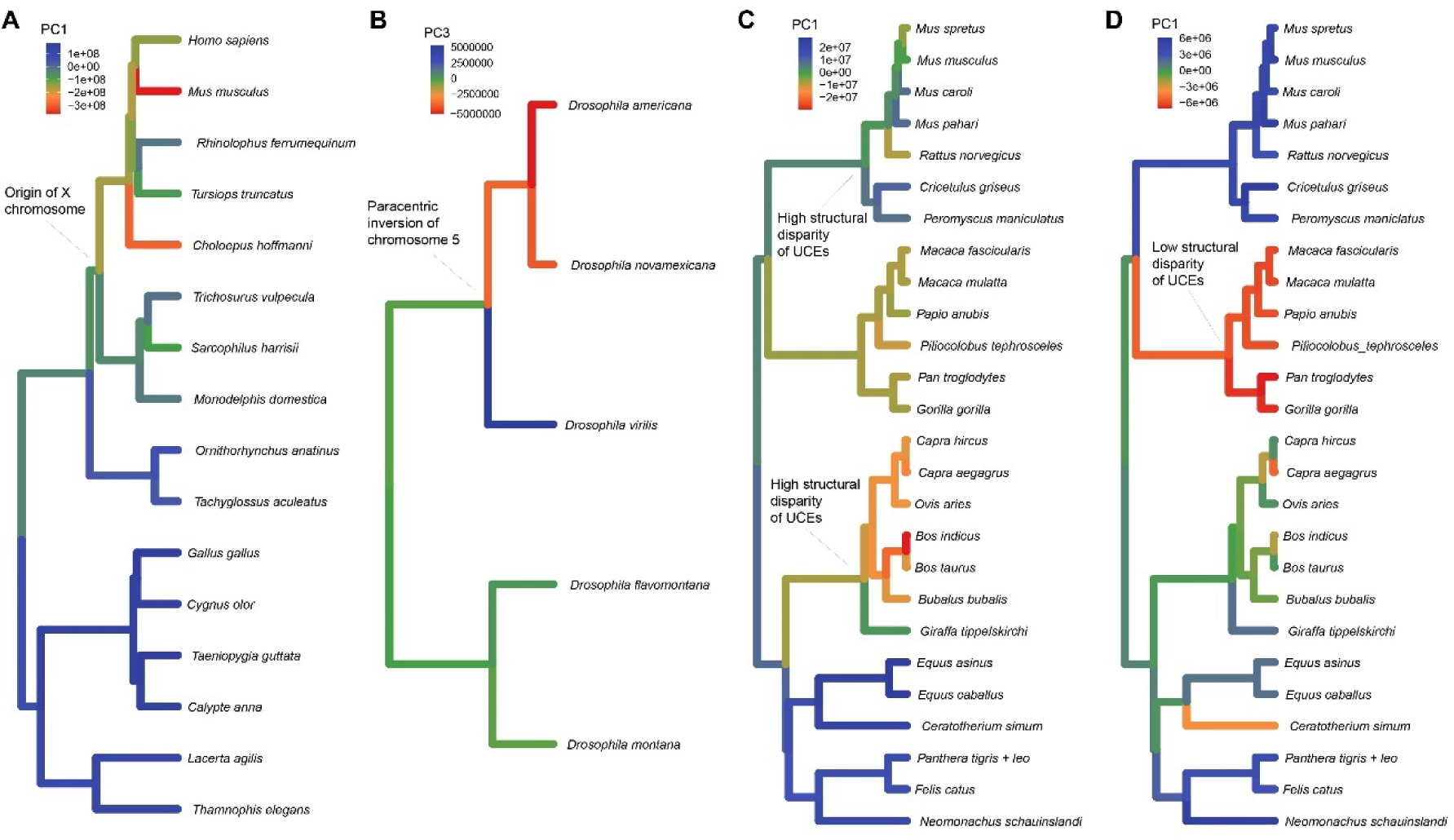
Ancestral state estimation of structural variants (SV) using select principal component (PC) scores from disparity analyses conducted herein. Structural variants detected using disparity analysis often have phylogenetic signal (Table 2) and can be used to quantitatively study the rate of SV evolution. Examples from this study include human X-linked BUSCO gene disparity of amniote vertebrates where structural disparity increases following the evolution of the X chromosome (PC1, A), the evolution of a speciation-associated paracentric inversion in chromosome 5 of the *Drosophila virilis* group that caused rearrangements in the position of UCE landmarks (PC3, B), and the identification of placental mammal orders with the least (chromosome set 1, PC1, C) and most (chromosome set 2, PC1, D) conserved UCE structure, respectively. Phylogenies were obtained from Kumar et al. [63].

**Table 3.** Results of phylogenetic signal tests using K [64] and λ [65] statistics for three most explanatory principal components (PCs) of structural variant disparity from the empirical analyses conducted herein. P-values are based on 1,000 randomizations (K) and likelihood ratio tests (λ). Significant tests are indicated with bold text.

| Principal component | Variance explained (%) | Phylogenetic signal |  |
| --- | --- | --- | --- |
| | | K | $\lambda$ |
| X-linked BUSCO landmarks from amniotes: PC1 | 71.2 | **1.3418 (p = 0.001)** | **1.160 (p < 0.001)** |
| X-linked BUSCO landmarks from amniotes: PC2 | 12.1 | 0.4237 (p = 0.501) | < 0.001 (p = 1.0) |
| X-linked BUSCO landmarks from amniotes: PC3 | 7.5 | **0.6591 (p = 0.049)** | 0.825 (p = 0.454) |
| UCE landmarks from *Drosophila*: PC1 | 94.0 | 0.6935 (p = 0.568) | **1.151 (p = 0.013)** |
| UCE landmarks from *Drosophila*: PC2 | 5.2 | **0.9949 (p = 0.027)** | **1.151 (p = 0.004)** |
| UCE landmarks from *Drosophila*: PC3 | 0.07 | 0.6831 (p = 0.195) | < 0.001 (p = 1.0) |
| UCE landmarks placental mammals (set 1): PC1 | 95.3 | **0.2173 (p = 0.022)** | **0.866 (p < 0.001)** |
| UCE landmarks placental mammals (set 1): PC2 | 4.1 | **0.2600 (p = 0.005)** | **0.818 (p < 0.001)** |
| UCE landmarks placental mammals (set 1): PC3 | 0.5 | 0.0643 (p = 0.717) | **0.476 (p = 0.004)** |
| UCE landmarks placental mammals (set 2): PC1 | 98.6 | **0.2677 (p = 0.009)** | **0.849 (p < 0.001)** |
| UCE landmarks placental mammals (set 2): PC2 | 0.8 | 0.1104 (p = 0.294) | **0.733 (p < 0.001)** |
| UCE landmarks placental mammals (set 2): PC3 | 0.5 | 0.0573 (p=0.708) | 0.659 (p = 0.235) |

## Discussion

Using a morphometrics-inspired approach we identified SVs associated with chromosome inversions, insertion-deletion events (=relative size change), and phylogeny. These qualities mirror the utility of morphological disparity analysis as applied to complex phenotypic structures. In a phenotypic context, disparity can be descriptive or serve as a proxy for phylogenetic affinities and ecology [22]. For example, disparity is widely used to understand adaptive radiation in the paleontological record [36]. Our simulation and empirical results suggest that, like classic morphological disparity, structural disparity of chromosomes increases through time (Fig. 7). In a genomic context, we are optimistic that disparity can play a similar role in understanding how chromosome architecture varies among species and can be used for quantitative comparisons to study organismal evolution and adaptation. Below we discuss the results of our empirical case studies, suggest guidelines for best practice when conducting structural disparity analysis, and present several future directions for applications and methods development.

Structural disparity and the evolution of chromosome architecture. Disparity analyses of SVs enable a broad understanding of how similar (or dissimilar) chromosome structures are across different species based on the distribution of conserved landmark sequences like BUSCO genes and UCEs. This analytical quality is particularly advantageous for studying large-scale shifts in chromosome organization across organisms relevant to the study of supergenes and homologous synteny blocks (sensu [16]). In our study we found that following the evolution of a key mammalian sex chromosome, structural disparity of 53 BUSCO genes increased compared to other amniotes (Fig. 2A, Fig. 8A). This disparity increase appears to be associated with an overall increase in chromosome size in those species where the landmark genes are located on an X chromosome. Due to the cessation of regular recombination, size increases and decreases in vertebrate sex chromosomes are well-documented [37]. When we scaled the positions by chromosome size, we observed that placental and marsupial mammals were no longer the most disparate group (Fig. S2E, G), further supporting chromosome size increase as the key driver of structural disparity among BUSCO genes in the mammalian X chromosome. Following size-correction, structural disparity analysis identified high disparity among the bird chromosomes analyzed, which from viewing the synteny plot (Fig. 2B) appears to be related to several inversions (within birds) associated with the landmarks. The ability for structural disparity analyses to capture SVs associated with inversions in closely related species or populations was also observed in our study of chromosome 5 from the *Drosophila virilis* group based on 191 UCE landmarks (Fig. 3). In this simple example, we observed that the first three principal components all corresponded to distinctive changes previously linked to speciation [27]. Although not all axes had statistically significant phylogenetic signal (Table 3), small sample sizes may have contributed to this outcome (see 38) as even those axes without significant signal (e.g. PC3) were consistent with phylogenetically associated changes (Fig. 8B).

The de novo study of structural disparity using tetrapod UCEs with placental mammals revealed order-specific disparity patterns in PCs 1 and 2 of both chromosome sets (Fig. 5). In addition to revealing clear phylogenetic patterns in UCE-related structural disparity, these analyses also revealed that disparity patterns are order-specific across different chromosomes (Fig. 8C, D). Specifically, using the chull function in the R package ‘sp’ [39] we found that convex hull values for each mammalian order on PCs 1 and 2 were perfectly correlated across chromosomes set 1 and set 2 (Pearson’s correlation coefficient = 0.996, p = 0.0003). This suggests that order-specific structural disparity associated with UCEs may be similar across different genomic regions. Given the importance of UCEs in development and gene regulation [40] these types of genome-wide patterns are important to explore as additional genomic data become available.

Best practices for structural disparity analysis. Based on comparative analyses of the X-linked BUSCO genes from amniote vertebrates and UCEs from placental mammals, we can recommend a set of best practices for structural disparity that will allow users to avoid several analytical artifacts that we encountered. First, examining component loadings for each landmark is important for identifying outlier landmarks that may obscure more generalized signal of other landmarks. In our study of placental mammals, we found that a single outlier (UCE 6948) explained most variation in the first two PCs (Fig. S6). The presence of this UCE in both datasets demonstrates that it is not single-copy and justifies removing these types of spurious landmarks prior to disparity analysis. Second, using raw, absolute positions may lead to outlier taxa if the landmarks are clustered at one end of a large chromosome. This phenomenon was observed in both the X-linked BUSCO gene dataset and the placental mammals (Fig. S2B, S7B, S8B). In some cases, scaling the positions based on chromosome size may resolve this issue (Fig. S2F), whereas in others it does not (Fig. S7F, S8F). We found that the use of bound landmark positions resolves this analytical artifact associated with the directionality of the chromosome sequence. If size is an important parameter for understanding structural disparity, we recommend using bound, absolute positions (Fig. S2C, S2D, S7C, S7D, S8C, S8D). If patterns like inversions and landmark translocations are of primary interest, we recommend using size-corrected and bound landmark positions (Fig. S2G, S2H, S7G, S7H, S8G, S8H). Finally, if using the de novo approach and clustering chromosomes with MMDS to identify homologous landmark regions for analysis we recommend viewing various axes as bivariate plots (Fig. 4B, 4C) where clustering may be more obvious than gap threshold sizes would suggest on the individual axes (Table 2).

A complementary tool for studying structural changes in chromosomes. Dimensionality reduction is already in wide use in comparative genomics (e.g., uniform manifold approximation and projection, [41]). Structural disparity analysis is therefore a natural companion method for assessing structural differences across the chromosomes of different species. It complements existing methods like WGA and synteny analysis by capturing a different dimension of how chromosomes vary across species. Synteny analysis is a powerful tool for identifying similarities in gene order and arrangement. While synteny analysis does capture spatial variation (unlike WGA), it is known to be very sensitive to parameter settings [42]. Furthermore, some methods of synteny inference are performed on post-WGA datasets meaning that structural information about the whole chromosomes has been removed a priori [15]. In contrast, landmark-based estimates of genome structure offer at least two qualities that syntenic analyses lack (Fig. S12). First, minor chromosomal rearrangements will be interpreted as breaks in synteny and ignored. In other words, inversions within syntenic regions not involving all the contiguous markers will break synteny. Second, large gap regions between contiguous orthologs can also break synteny (depending on parameter settings). The latter example highlights another important difference; disparity analyses will capture distance information between landmarks whereas synteny only provides landmark (ortholog) order information.

Applications beyond diploid genomes and evolutionary biology. Apart from diploid organisms such as vertebrates and insects, the structural disparity pipeline could also be implemented on polyploid organisms when chromosome phasing has been performed. Utilizing the different phased genome assemblies, the pipeline can identify structural differences between haplotigs of the genome. In addition, the pipeline may be useful for measuring structural evolution of pangenomes without the need for a reference input genome. Instead, a class/category of genetic markers could be used as structural landmarks. As a visualization tool, structural disparity analysis may complement the routinely used pangenome graph building methods (e.g.[43]). Lastly, a potential use of structural disparity analysis relates to cancer genomics. The genomes of somatic cancer cells undergo rapid structural evolution and SVs in cancer cells are very specific to the type of cancer [44]. SVs in cancer cells have an impact on gene expression through modifying structural accessibility, DNA methylation (and more) in the cis-regulatory regions of the genome [44], therefore the ability to identify genomic rearrangement is valuable (especially when using long-read data).

Avenues to expand the method towards genomic disparity. Structural disparity analyses could be developed for whole-genome comparisons. At present, the pipeline performs chromosome to chromosome comparisons, limiting comparisons across genomic regions that have undergone translocation, fission, or fusion events. With greater awareness of the frequency of these events, weighting parameters could be defined to measure global genomic disparity levels. While we limited our validation analyses to using BUSCOs and UCEs as landmarks and a handful of (largely homologous) chromosomes, landmarks could be any conserved genetic sequences which can be accurately mapped. Variation in the placement of conserved sequences can be used to study colocalization of transcribed genes (e.g. BUSCO [45]), cis-regulatory elements, enhancers, promoting regions, and more. The ethos of the approach is also scalable to three-dimensional genome structure once that information is more broadly available for many species (see [46, 47]). The extensive literature on geometric morphometrics offers substantial guidance for how 3D comparisons may be performed in the future [25]. Landmark-based estimates of genomic disparity provide a rapid mechanism for comparisons of genome structure across species and populations, and we anticipate they will expedite discovery and study of novel structural patterns within comparative genomics.

## Methods

Bioinformatics and identification of landmarks. The bioinformatics and computational analyses were performed on Crop Diversity HPC, described by Percival-Alwyn et al. [48]. The structural disparity pipelines are modular and can be modified for a variety of other uses. Three step-by-step tutorials along with custom R and shell scripts are available on GitHub (https://github.com/nhm-herpetology/genomic-disparity/blob/main/README.md). Structural disparity analysis can be conducted using a variety of genomic landmarks assuming they are single copy and orthologous (e.g. BUSCO genes [45], UCEs [32,34]). In this study, we used human X-linked BUSCO genes and UCEs as landmarks. We selected human X-linked genes because many of them are found in homologous blocks on a single chromosome across amniote vertebrate animals [18] and UCEs because they are highly conserved, lack repeats, and widely distributed in genomes [34, 49]. UCEs have also been implicated as relevant to the structure of key developmental and transcriptomic pathways [49, 50], making variation in their relative chromosomal placements of biological interest.

Sensitivity analyses related to chromosome size, complementarity and landmark boundaries. Data transformations associated with object size are a well-studied component of morphometrics [51]. In our datasets where structural disparity was correlated with chromosome size, we conducted a series of analyses to establish best practice for structural disparity analysis. We analyzed these datasets in four ways using: (1) raw positions, (2) bounded, raw positions, (3) chromosome size-corrected positions, and (4) bounded, size-corrected positions. We repeated each analysis using the reverse complement version of the four datasets for a total of eight versions of the analysis. Boundary setting was achieved for each landmarked chromosome using a two-step procedure: (1) re-positioning all the landmarks relative to the first landmark and (2) using the position of the last landmark to establish the size of the genomic region being analyzed. Absolute size correction was performed with R functions that divided the total chromosome size by each landmark position. Bounded size correction was achieved by dividing the size of the bounded genomic region by each landmark position with R functions.

BUSCO genes and amniote vertebrates. Using the BUSCO gene annotation file from Lovell et al. [18], we extracted all the genes annotated in the human (*Homo sapiens*) X chromosome. We utilized this list of x-linked genes and identified homologous chromosomes in other species based on the proportion of the human x-linked genes mapped on them. The other species included in the study were four placental mammals (*Mus musculus*, *Rhinolophus ferrumequinum*, *Tursiops truncatus*, and *Choloepus hoffmanni*), three marsupial mammals (*Sarcophilus harrisii*, *Monodelphis domestica*, and *Trichosurus vulpecula*), two monotreme mammals (*Tachyglossus aculeatus* and *Ornithorhynchus anatinus*), four birds (*Gallus gallus*, *Cygnus olor*, *Calypte anna*, and *Taeniopygia guttata*) and two squamate reptiles (*Thamnophis elegans* and *Lacerta agilis*). When there was more than a single chromosome that shared human x-linked genes (e.g. *Tachyglossus, Ornithorhynchus*, and *Trichosurus*), the chromosome with the higher proportion of genes was used to maximize the number of landmarks available for structural disparity analysis. Lovell et al. (2022) also analyzed the budgie (*Melopsittacus undulatus*) genome, however, because our pipeline did not identify focal landmarks on a single chromosome, we did not include this species in the structural disparity analysis. Using the gff3 file approach (Fig. 1), we identified human x-linked BUSCO genes that were homologous across all the species’ chromosomes.

Detailed instructions for identifying and extracting these genes are available in Step 1 of the GitHub tutorial. Next, chromosomes are prepared for downstream analysis using a pipeline which requires the following R statistical software [31] packages: rehshape2 [52], matrixStats [53], tidyr [54], dplyr [55], and ggplot2 [56]). From 838 genes we identified 53 that were present in all focal species making them suitable for use as landmarks in a structural disparity analysis. Detailed instructions for identifying and extracting landmarks are available in Step 2 of the GitHub tutorial. We also conducted data matrix manipulation and principal component analysis in R, with code available in Step 3 of the GitHub tutorial. For size-corrections, we obtained chromosome sizes from NCBI. We tested for correlations between chromosome size and disparity using Spearman’s rho nonparametric tests in R. To create a synteny plot for comparative purposes, we used the Chromomap package in R [35]. A list of the 53 human X-linked BUSCO genes used as landmarks can be found in Appendix S1.

Drosophila structural variants, speciation and UCEs. Based on the findings of Poikela et al. [27], we landmarked chromosome 5 for *D. americana*, *D. flavomontana*, *D. montana*, *D. novamexicana*, and two individuals of *D. virilis*. This chromosome has known inversions and translocations and may have contributed to speciation in the group. We downloaded chromosome 5 from the *Drosophila virilis* group either via NCBI Entrez Direct UNIX E-utilities (2) or directly from the literature (1). The following chromosomes sequence were used: *D. americana* (NCBI accession no: CM061086.1), *D. flavomontana* (1), *D. montana* (1), *D. novamexicana* (CM061080.1), and two individuals of *D. virilis* (CM017608.2 and CM061075.1). Detailed instructions for downloading chromosome sequences are available in Step 1 of the GitHub tutorial. For reference mapping of ultraconserved elements (UCEs), we utilized a published set of ultraconserved elements (UCEs) from dipterans [3]. We indexed the chromosome assemblies and reference mapped UCEs using default settings in BWA-MEM [29] and SAMtools [57]. Custom shell commands are used to loop the reference mapping and create a spreadsheet that contains presence and absence information for the UCEs on each chromosome 5 variant included in the analyses. Detailed instructions on this reference mapping approach are available in Step 2 of the associated GitHub tutorial. Next, chromosomes were prepared for downstream analysis using an R pipeline as described above. The pipeline includes three steps (i) remove mapped UCEs that the chromosomes do not share, (ii) complementarity checks to ensure all chromosomes have comparable orientations (5’ to 3’), and (iii) merging redundant UCE probes that target the same UCE locus. The pipeline resulted in the identification of 191 homologous UCE landmarks for chromosome 5 of the *Drosophila virilis* group. Detailed instructions for the preparation pipeline are available in Step 3 of the GitHub tutorial. In the final step of the disparity analysis, we conducted a principal component analysis (PCA) and visualized the results. The input file for the PCA is available on GitHub as ‘Drosophila_PCA.csv’. We used R base functions along with dplyr to format the data matrix and conduct the PCA. Visualization was conducted using ggplot2 [56] and cowplot [58]. Detailed instructions for this final step are available in Step 4 of the GitHub tutorial.

De novo landmark discovery and placental mammal genomes. To apply our disparity pipeline to a group where homology among chromosomes was unknown, we used 26 nuclear genomes from five orders of placental mammals (Table 1). Because placental mammal orders have diverged over millions of years [59] the homology of their chromosomes is fragmentary, making them a good candidate for exploring the extent to which disparity analysis can be applied to single genomic regions in divergent groups. We downloaded mammal genome assemblies using a custom script (chromosome_download.sh) that we developed to work with a standard input file (chromosome_download.config). Setting up a batch download for a subset of the mammal taxa (26 species) is described in detail in Step 1 of the GitHub tutorial. For reference mapping, we utilized a published set of tetrapod UCEs [32]. We indexed and reference-mapped UCEs on the 1,004 mammal chromosomes using default settings in BWA-MEM [29] and SAMtools [57]. Reference mapping was conducted using a custom shell script that loops through the chromosome sequences (landmark_mapping.sh). We tested the utility of minimap2 [30] for mapping conserved regions to the genome assemblies utilizing default settings on chromosome set 2 of the mammalian genomes. We then compared these results to those obtained using BWA MEM. We observed that BWA MEM mapped higher numbers of UCEs than minimap2 (Fig. S1). This difference in mapping quality could be explained by the short length of the UCE sequences used, at about 180 bp length [32] because BWA MEM can achieve higher accuracy with short-read mapping and produces less false negatives than minimap2 [30]. However, minimap2 may be preferable if larger sequences were used as landmarks, and that approach could then be easily combined with our post-mapping pipeline steps. In addition to providing a UCE presence-absence output file for downstream analysis in the pipeline, landmark_mapping.sh outputs the number of (UCE) landmarks mapped and length information for each chromosome. Detailed instructions for refence mapping are available in Step 2 of the GitHub tutorial.

Interestingly, we found that the number of UCEs mapped onto each mammal chromosome was positively and significantly correlated with chromosome size (r = 0.75, p < 0.05; SI Appendix, Fig. S2). To identify sets of chromosomes that share homologous landmarks, we applied metric multidimensional scaling (MMDS) on the presence-absence matrix generated by landmark_mapping.sh. Using MMDS scores across 10 axes we identified mutually exclusive sets of chromosomes that consisted of only one chromosome from a species. These chromosome sets were identified by selecting a threshold value on the relevant axes of the MMDS plot. Detailed instructions for clustering chromosomes and thresholding are available in Step 3 of the GitHub tutorial. To determine if protein content was similar on chromosomes sets (implying that they share functional, protein-coding homologs in addition to UCE landmarks, we used protein annotations from the Ensembl database (release 110; [60]), to generate Jaccard similarity coefficients for comparing protein compositions on the two focal chromosome sets across a subset of placental species where annotations were available (N = 14). After extracting the protein-coding markers’ identities from the Ensembl annotation files of each species (for the specific chromosomes in set 1 and set 2 separately), we utilized a custom defined Jaccard’s similarity index which removed case sensitivity in the names of the markers (available on GitHub page as ‘Jaccard_proteins.R’). This index estimated pairwise similarities between each chromosome in set 1 and set 2 respectively, as well as for all chromosome pairs in both the sets. The latter was done to test if chromosomes within a set show higher similarity in protein-coding markers than chromosomes between the two sets. Following the identification of chromosome sets, we prepared them for PCA by (i) removing UCEs that did not occur on all chromosomes, (ii) checking the directional orientation (5’ to 3’), and (iii) merging redundant UCE probes. Unlike the highly conserved chromosomes from the Drosophila example, determining the directionality of mammal chromosomes was not always obvious. In other words, sometimes we observed mixed forward and reverse mappings of UCE landmarks, suggesting that only particular segments of the landmarked region were sequenced in the opposite direction of most taxa. This is likely explained by within-chromosome recombination and to account for it we used a majority rule criteria so that we only changed chromosome orientations if >50% of landmarks were mapped in the opposite direction of the reference chromosomes. Detailed instructions for the chromosome preparation pipeline are available in Step 4 of the GitHub tutorial. Finally, we conducted a PCA on each of the chromosome sets and visualized the results which are detailed in Step 5 of the GitHub tutorial. The input files for the PCA analysis are available on GitHub as ‘Set_1_mammals_PCA.csv’ and ‘Set_2_mammals_PCA.csv’, respectively. As with the Drosophila example, applying the disparity pipeline to placental mammal genomes required the following R packages: rehshape2, matrixStats, dplyr, ggplot2 and cowplot.

Simulation and other tests for increased disparity through time. We simulated two common evolutionary shifts in chromosome structure that occur as species and populations diverge: insertion-deletion (indel) events and chromosomal inversions. We performed these simulations using chromosome set 1 from three species subsampled from the larger dataset, *Capra aegagrus*, *Mus pahari* and *Neomonachus schauinslandi* (Table 1). We selected these taxa based on phylogenetic diversity (representing Artiodactyla, Carnivora and Rodentia) and being outliers in some analyses (*Capra aegragrus*, Fig. S9). We simulated evolutionary events using SimuG [61]. We designed the experiment logarithmically, simulating 1, 10, 100, 1,000, and 10,000 random indels or inversions on the ancestral chromosome of the five focal species. We ran five replicated simulations for each of the five focal species. To assess the change in disparity between ancestral chromosomes and their simulated descendants, we used a simple pairwise metric of increase in disparity (ID):

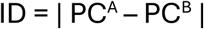

Where PC^A^ is the principal component score of the ancestral chromosome, PC^B^ is the principal component score of the descendent chromosome, and ID is calculated by taking the absolute difference of these values. As such, ID is an intuitive measurement of how much landmark placement changed during the simulations, where a value of 0 indicates no change and larger values correspond to increasingly disparate landmark placements. We predicted that if genomic disparity reflects the degree of evolutionary change of architecture between two genomes, there should be a positive correlation between ID and the number of simulated events. While analysis of individual PCs may be problematic when data have high effective dimensionality [62], here the dominance of PC1 and the overwhelming correlation with chromosome size on that primary axis, justifies separate analysis of these components. We compared the original chromosome (set 1) sequence from three species of mammals, referred to as the ‘ancestral chromosome’, to simulated ‘descendent chromosomes’ which we made by simulating random indel or inversion events (see main text for more detail). Following the disparity pipeline described above, we mapped UCEs onto the simulated chromosomes, extracted homologous landmarks and carried out a PCA. PCA also included ancestral chromosomes from each species. We calculated the disparity between the ancestral chromosomes and simulated descendent chromosomes using the equation described in the main manuscript. Disparity was calculated using the logarithm of the absolute difference between the PC values of the ancestral chromosome and each simulated chromosome. Furthermore, we tested for differences between the disparity measures of indel-simulated and inversion-simulated chromosomes using two-way ANOVAs implemented in R.

To provide empirical context for the simulation results, we tested for the signature of disparity increase in our placental mammal datasets using nonparametric Wilcoxon tests in R. If UCE-inferred disparity increases with evolutionary time, we expected inter-order disparity to be significantly higher than intra-order disparity for the most explanatory PCs because there has been more evolutionary divergence between mammalian orders than within them. We also generated Jaccard similarity coefficients for comparing protein compositions between and within mammalian orders. Tests for phylogenetic signal and ancestral state estimation. Using the three most explanatory structural disparity PCs, we tested for phylogenetic signal in the (1) amniote vertebrate, (2) *Drosophila virilis* group, and (3) placental mammal (both chromosome sets 1 and 2) datasets. Phylogenies for each dataset were obtained from Kumar et al. [63]. We used two common metrics for phylogenetic signal, K [64] and λ [65]. We implemented test using the phytools package [66] in R. We assessed the significance of the tests using 1,000 randomizations (K) and likelihood ratio tests of the observed data to that estimated under Brownian motion (λ). To place structural disparity of genomic regions in a phylogenetic context, we estimated putative ancestral disparity values we used the ‘fastAnc’ function in phytools for select PCs in each dataset.

## Acknowledgments

We thank members of the Goswami Lab and NHM Herpetology Group for many helpful comments on experimental design and methods development. The authors acknowledge ChatGPT for assistance with some R coding (https://openai.com/index/chatgpt/). We thank Kate Thomas for an introduction to several R packages. Several delegates at the 2023 Society for Integrative and Comparative Biology (Austin, Texas, USA) and 2023 Evolution (Albuquerque, New Mexico, USA) and Young Investigators meeting 2024 (Bhopal, India) conferences provided valuable suggestions on earlier version of this work and putative applications for structural disparity analysis. We thank Jason Head and members of the Department of Zoology at the University of Cambridge for helpful suggestions related to size correction. We thank Christophe Liedtke for discussions on dimensionality reduction in genomics. We are indebted to Brant Faircloth and two anonymous reviewers for comments that greatly improved the quality of this manuscript. We are grateful to Iain Milne for bioinformatics support. The authors acknowledge Research Computing at the James Hutton Institute for providing computational resources and technical support for the “UK’s Crop Diversity Bioinformatics HPC” (BBSRC grants BB/S019669/1 and BB/X019683/1), use of which has contributed to the results reported within this paper. This study was part of a project funded by the European Commission through Marie Skłodowska-Curie Actions to A.V.M. (Grant Agreement ID: 101027832). This work also received support from the Genomics Research Theme at the Natural History Museum, London.

## Supplementary Information

**Figure S1.**
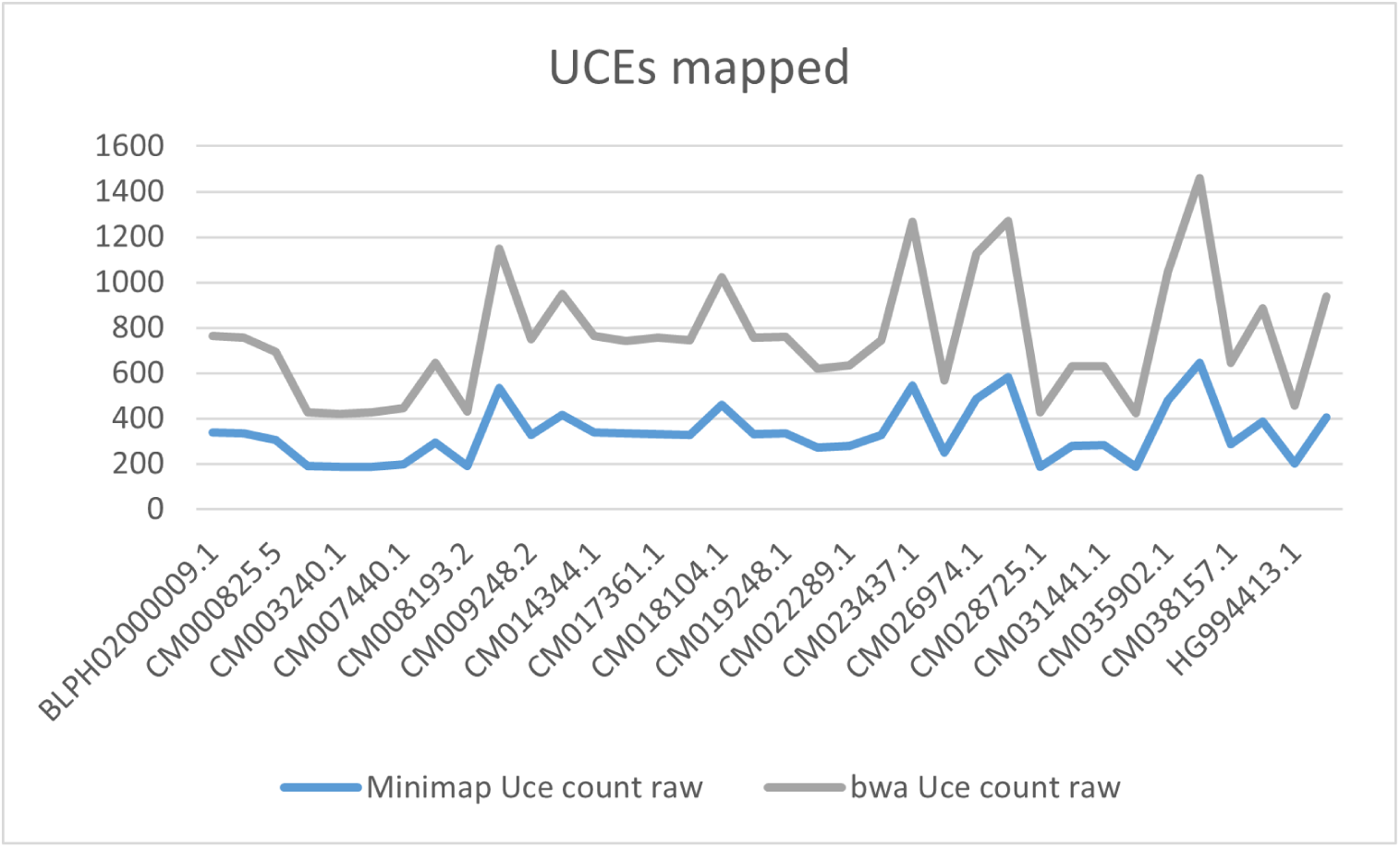
Number of ultraconserved elements (UCEs) mapped utilizing two different mapping software: minimap2 (blue) and bwa-mem (grey). Demonstrated using a subset of chromosome assemblies picked from the ‘chromosome set 2’ of the placental mammals UCE dataset discussed in the main text.

**Figure S2.**
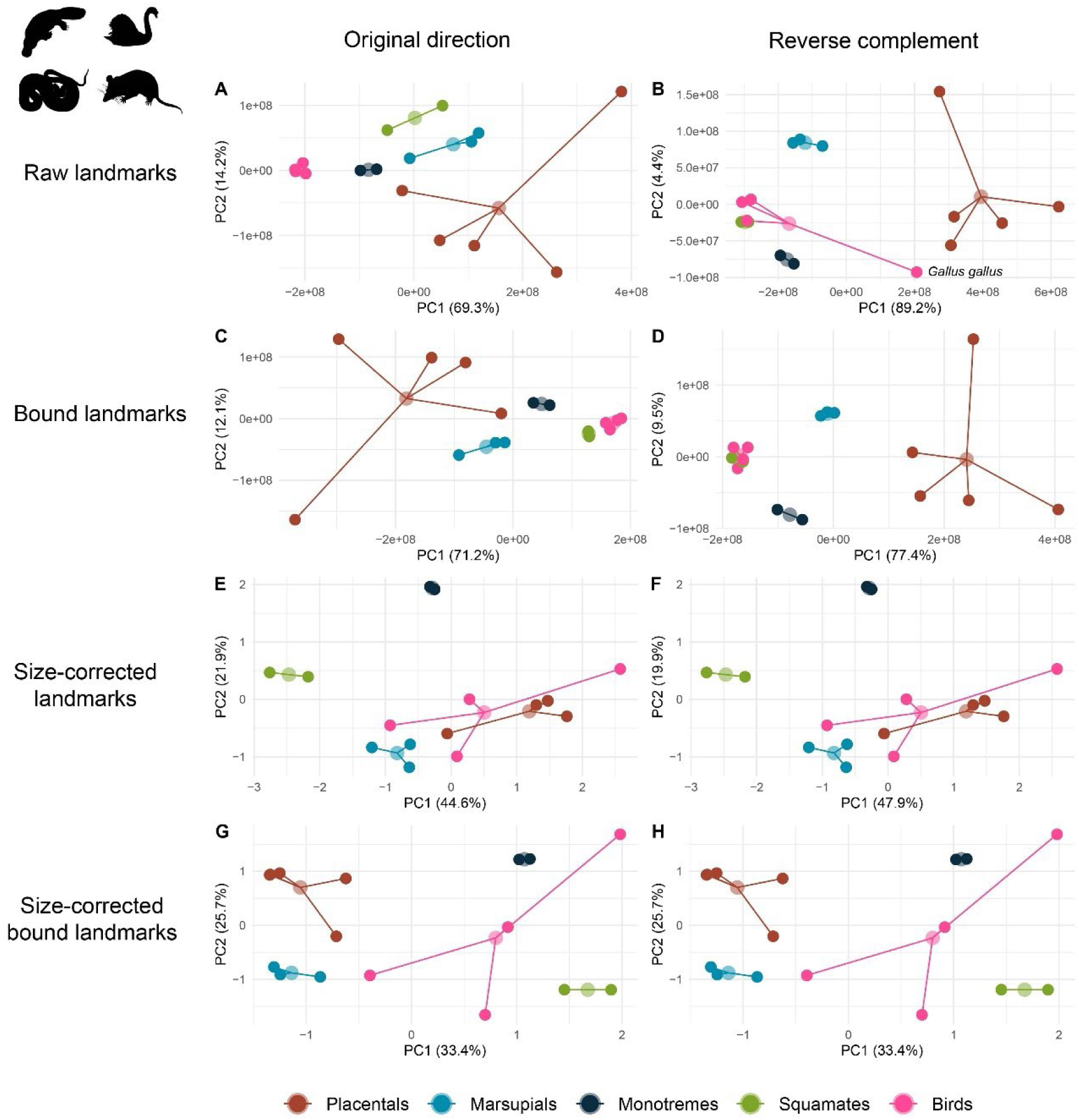
Principal component (PC) sensitivity analyses of 53 X-linked BUSCO landmarks in amniote vertebrates taken from Lovell et al. (2022). The first row are PCAs conducted using raw, absolute positions for the original direction (A) and reverse complement (B), respectively. The second row are PCAs conducted using bounded, absolute positions for the original direction (C) and the reverse complement (D). The third row are PCAs conducted using size-correction with chromosome size for the original direction (E) and the reverse complement (F). The fourth row of plots are PCAs conducted using size-correction using bounded chromosome sizes and positions for the original direction (G) and reverse complement (H). The chicken (*Gallus gallus*) is highlighted in plot B to illustrate an analytical artifact associated with clustered landmarks on large chromosomes. Organism silhouettes downloaded from https://www.phylopic.org/.

**Figure S3.**
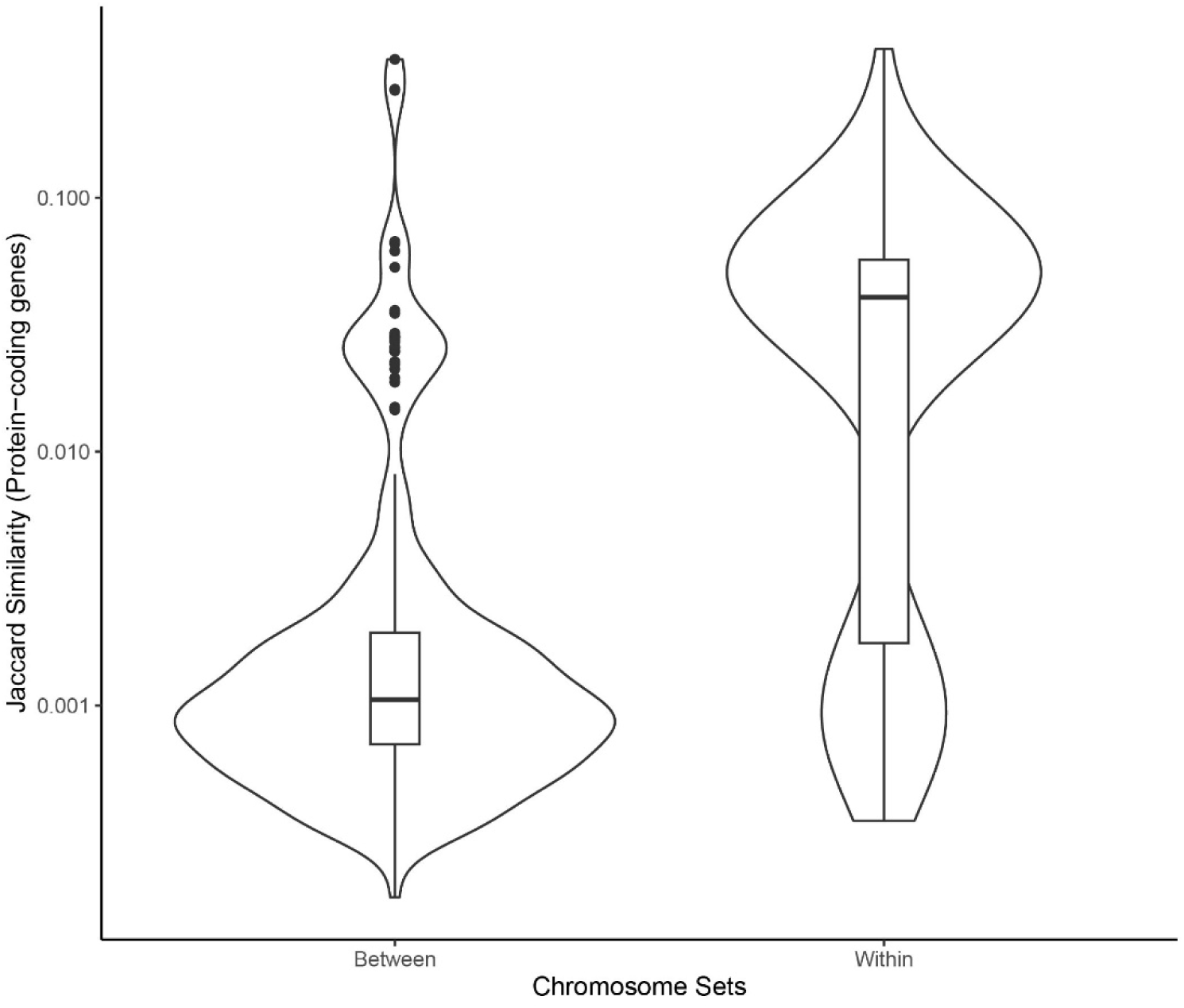
Jaccard similarity coefficients for within and between protein-coding gene content in the two focal chromosome sets used in the study.

**Figure S4.**
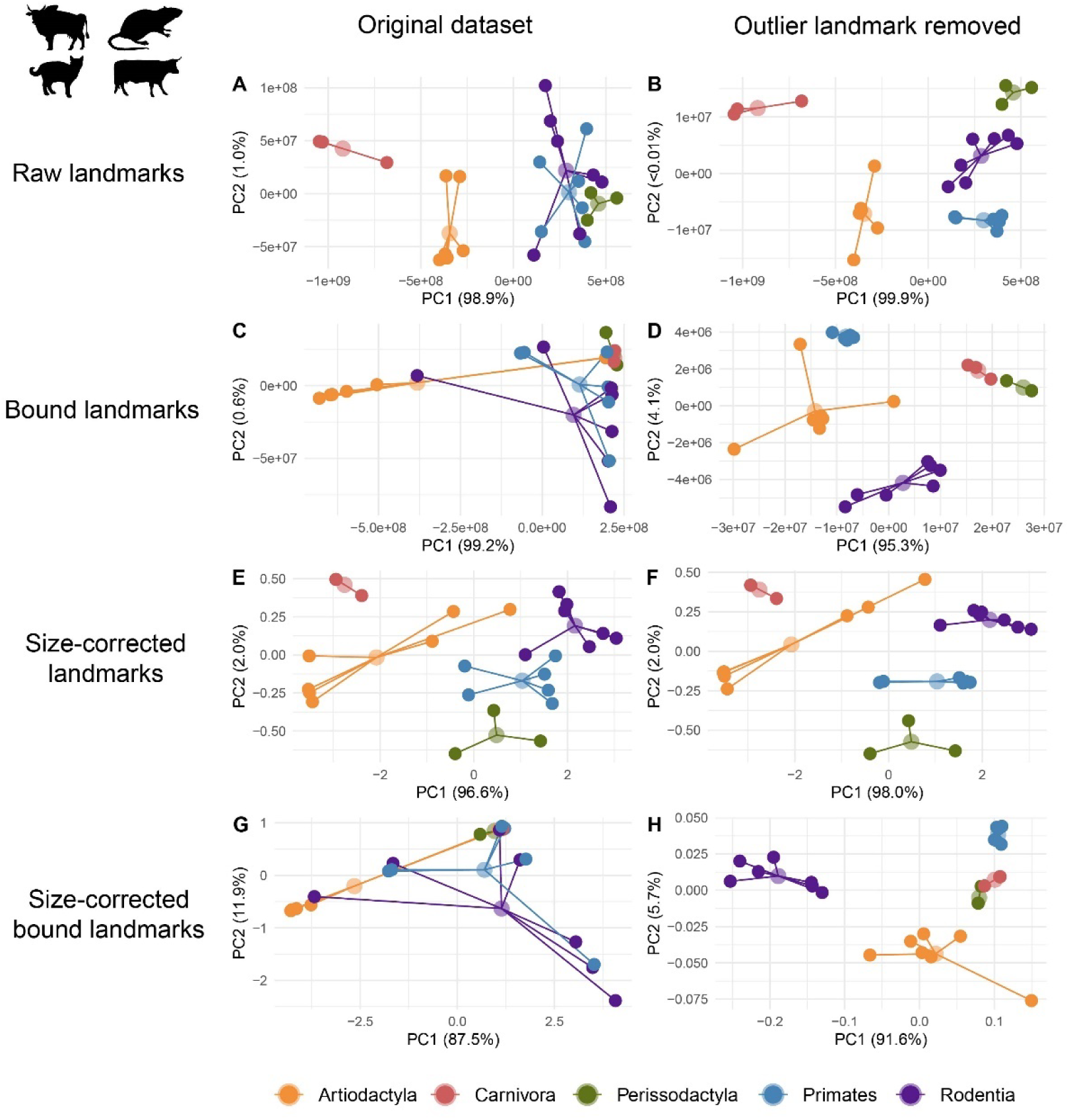
Principal component analyses (PCAs) of ultraconserved element (UCE) landmarks in placental mammals for chromosome set 1 before and after the removal of an outlier landmark (UCE 6948). The first row of plots are PCAs conducted using raw, absolute positions for the original direction (A) and reverse complement (B), respectively. The second row of plots are PCAs conducted using bounded, absolute positions for the original direction (C) and the reverse complement (D). The third row of plots are PCAs conducted using size-correction with chromosome size for the original direction (E) and the reverse complement (F). The fourth row of plots are PCAs conducted using size-correction using bounded chromosome sizes and positions for the original direction (G) and reverse complement (H). The first column of plots are PCAs conducted with all 186 UCE landmarks and the second column of plots are PCAs based on 185 UCE landmarks (outlier removed).

**Figure S5.**
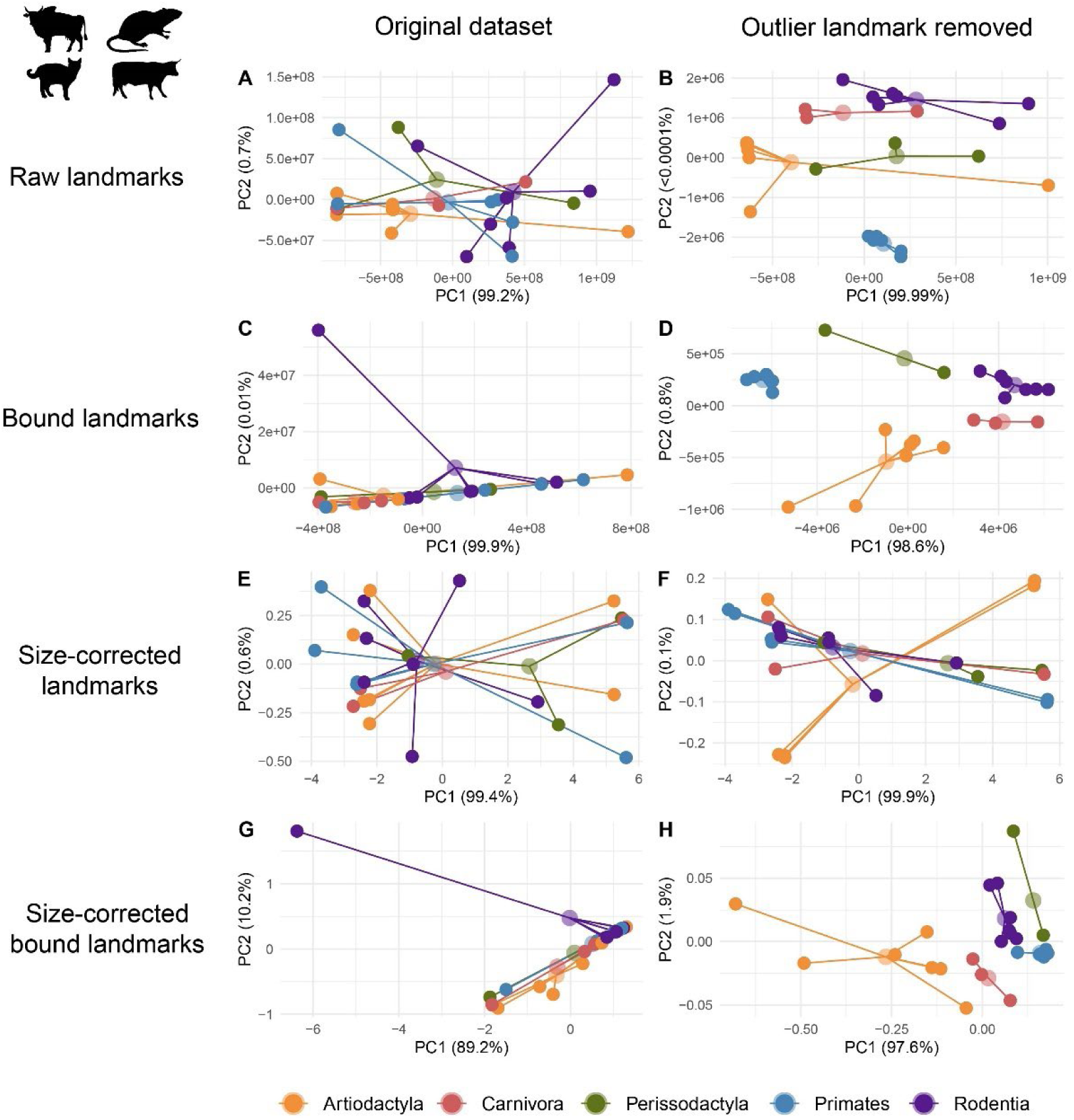
Principal component analyses (PCAs) of ultraconserved element (UCE) landmarks in placental mammals for chromosome set 2 before and after the removal of an outlier landmark (UCE 6948). The first row of plots are PCAs conducted using raw, absolute positions for the original direction (A) and reverse complement (B), respectively. The second row of plots are PCAs conducted using bounded, absolute positions for the original direction (C) and the reverse complement (D). The third row of plots are PCAs conducted using size-correction with chromosome size for the original direction (E) and the reverse complement (F). The fourth row of plots are PCAs conducted using size-correction using bounded chromosome sizes and positions for the original direction (G) and reverse complement (H). The first column of plots are PCAs conducted with all 82 UCE landmarks and the second column of plots are PCAs based on 81 UCE landmarks (outlier removed).

**Figure S6.**
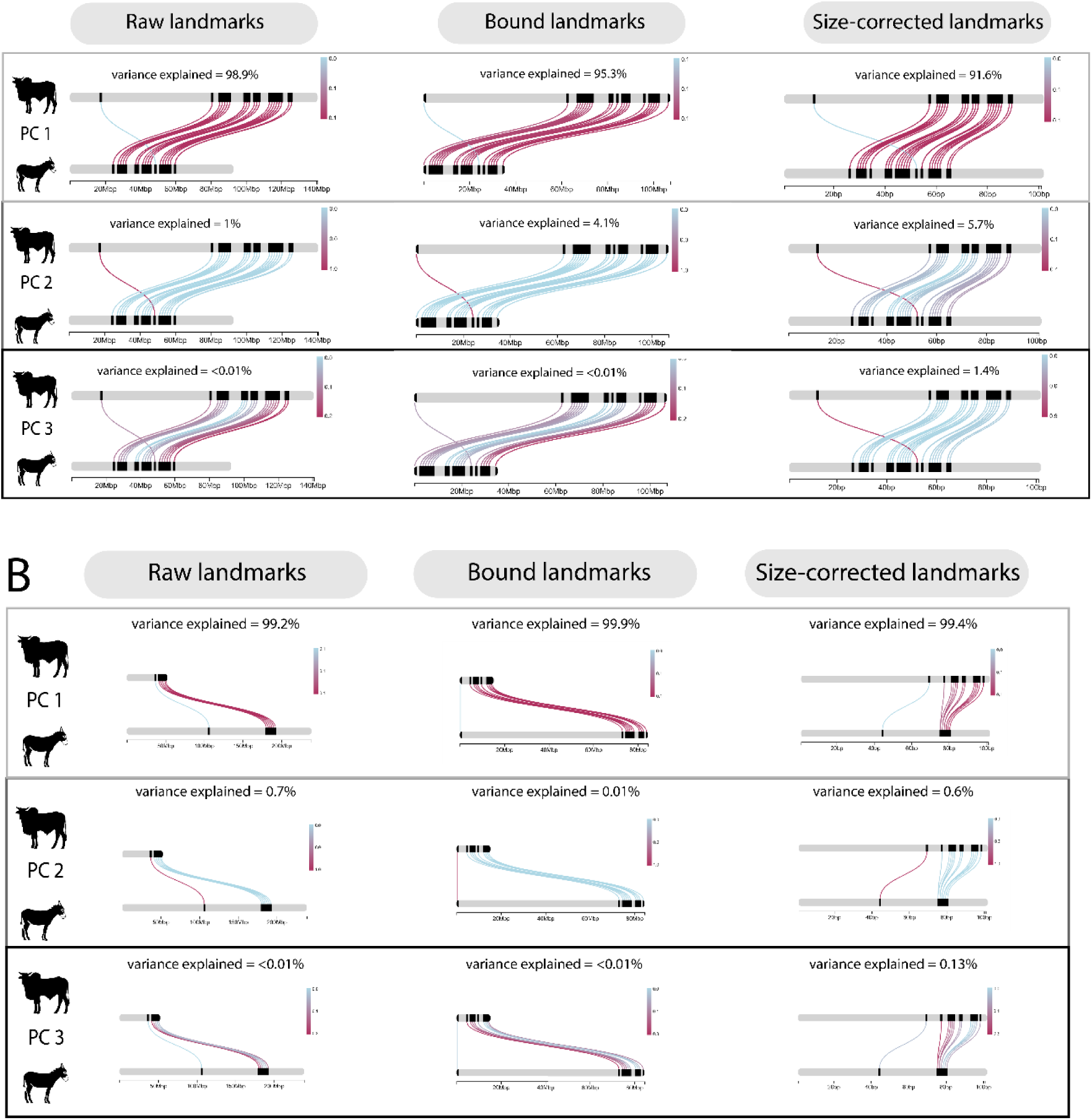
ChromoMap plots visualizing principal component analysis (PCA) loadings of landmarks on PC1, PC2, and PC3 in the raw landmark PCA, bounded landmark PCA, and size corrected landmark PCA of chromosome set 1 (A) and set 2 (B) from the placental mammal visualized using links between *Bos indicus* and *Equus asinus*. Respective variances explained by each principal component are listed in the plot. A single landmark (UCE 6948) was found to be uniquely weighed on the various PCs identifying it as an outlier landmark.

**Figure S7.**
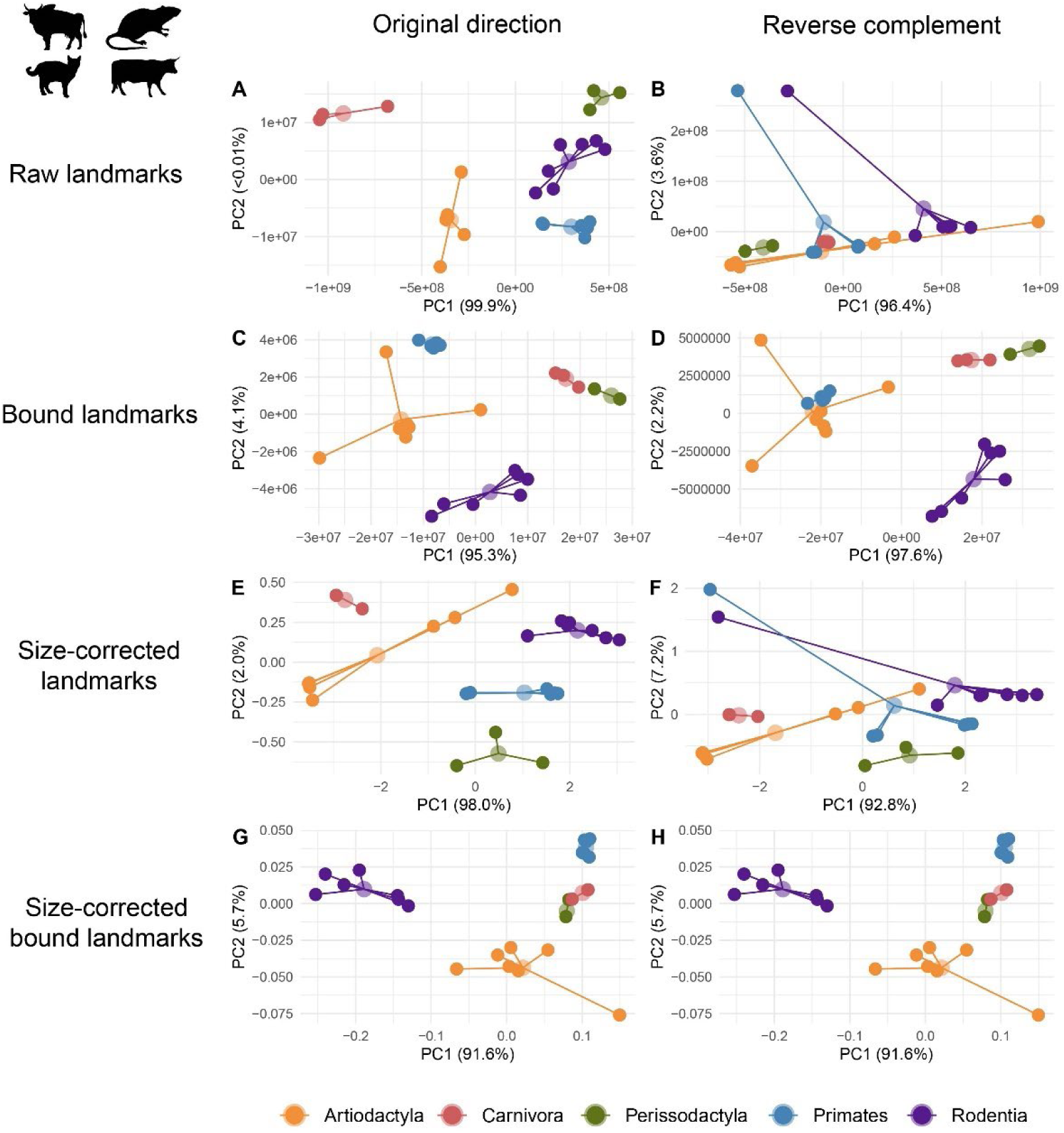
Principal component (PC) sensitivity analyses of outlier removed UCE landmarks in placental mammals chromosome set 1. The first row are PCAs conducted using raw, absolute positions for the original direction (A) and reverse complement (B), respectively. The second row are PCAs conducted using bounded, absolute positions for the original direction (C) and the reverse complement (D). The third row are PCAs conducted using size-correction with chromosome size for the original direction (E) and the reverse complement (F). The fourth row are PCAs conducted using size-correction using bounded chromosome sizes and positions for the original direction (G) and reverse complement (H).

**Figure S8.**
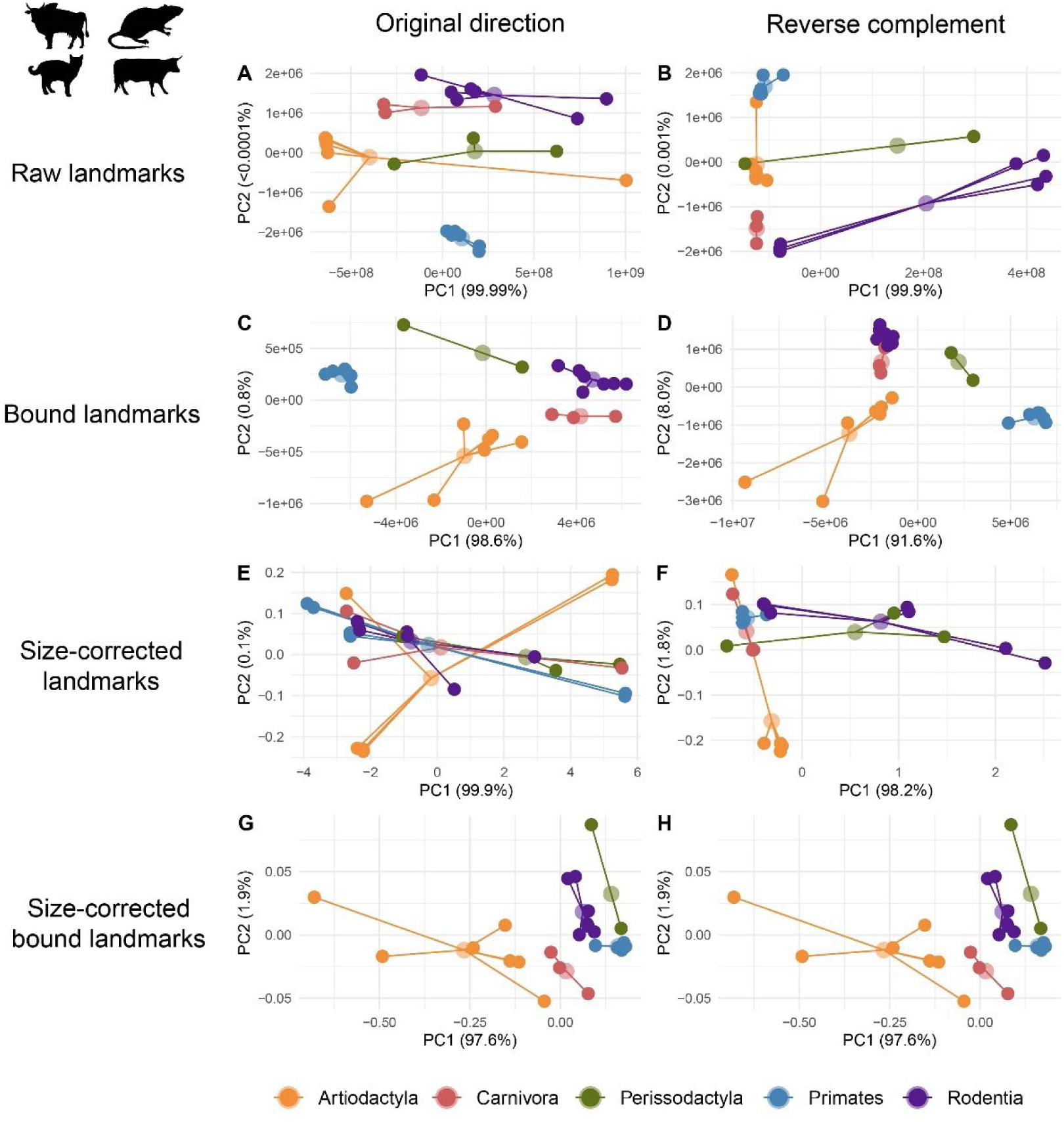
Principal component (PC) sensitivity analyses of outlier removed UCE landmarks in placental mammals chromosome set 2. The first row are PCAs conducted using raw, absolute positions for the original direction (A) and reverse complement (B), respectively. The second row are PCAs conducted using bounded, absolute positions for the original direction (C) and the reverse complement (D). The third row are PCAs conducted using size-correction with chromosome size for the original direction (E) and the reverse complement (F). The fourth row are PCAs conducted using size-correction using bounded chromosome sizes and positions for the original direction (G) and reverse complement (H).

**Figure S9.**
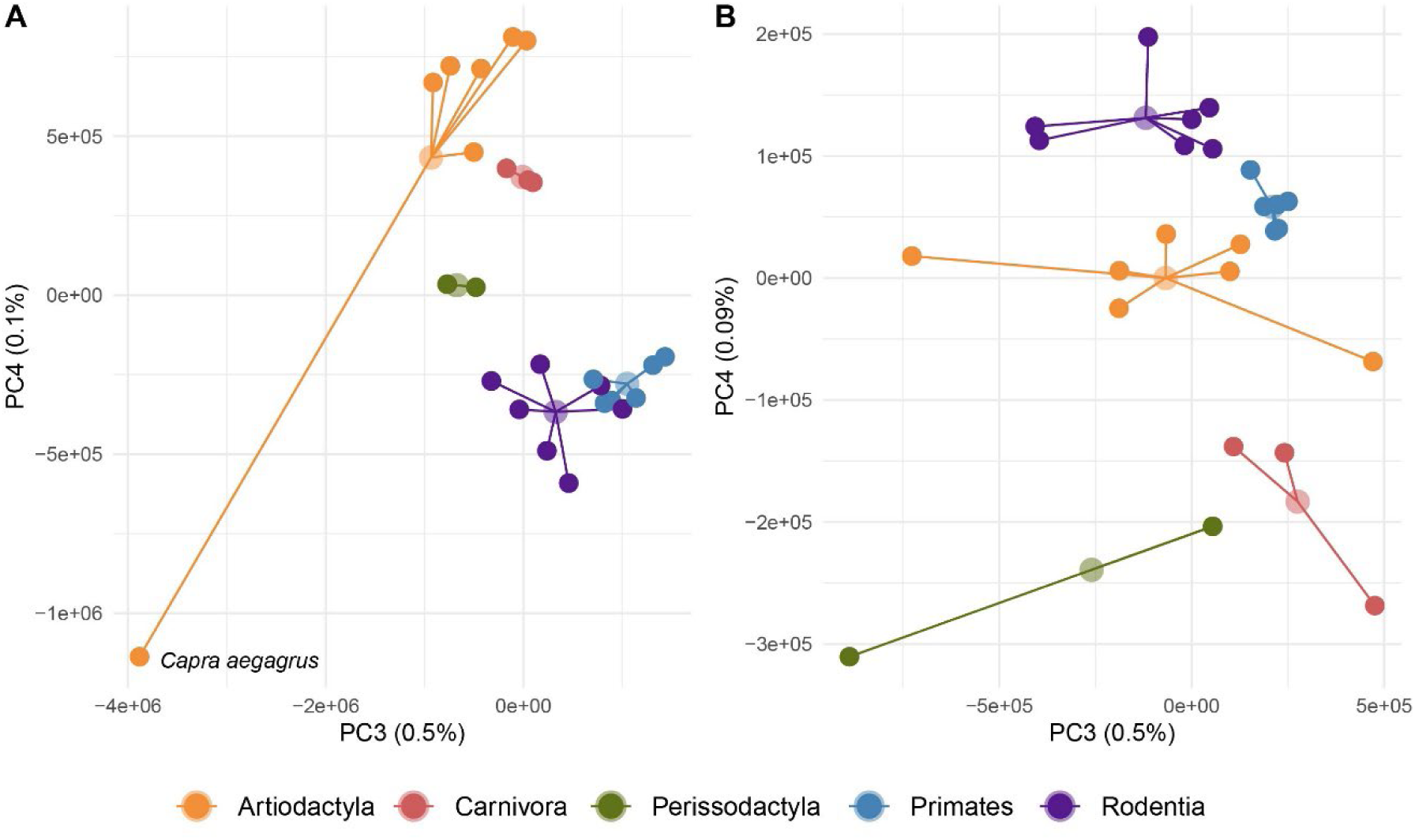
Ordination results from principal component (PC) analysis of UCE landmark positions in two sets of chromosomes from 26 species of placental mammals using bounded raw positions after the removal of an outlier landmark (see text). Representatives for five orders (Artiodactyla, Carnivora, Perissodactyla, Primates and Rodents) were included (Table1). The depicted structural variation is inferred from PCs 3 and 4 of 185 (chromosome set 1, A) and 81 (chromosome set 2, B) landmarks, respectively. In both datasets, the first two PCs explained > 99% of the variation (see **Figure 5**, main text). While explaining < 1% of the overall variation, PCs 3 and 4 mostly separated mammalian orders in geno-metric space. *Carpa aegragus* was a notable outlier in chromosome set 1.

**Figure S10.**
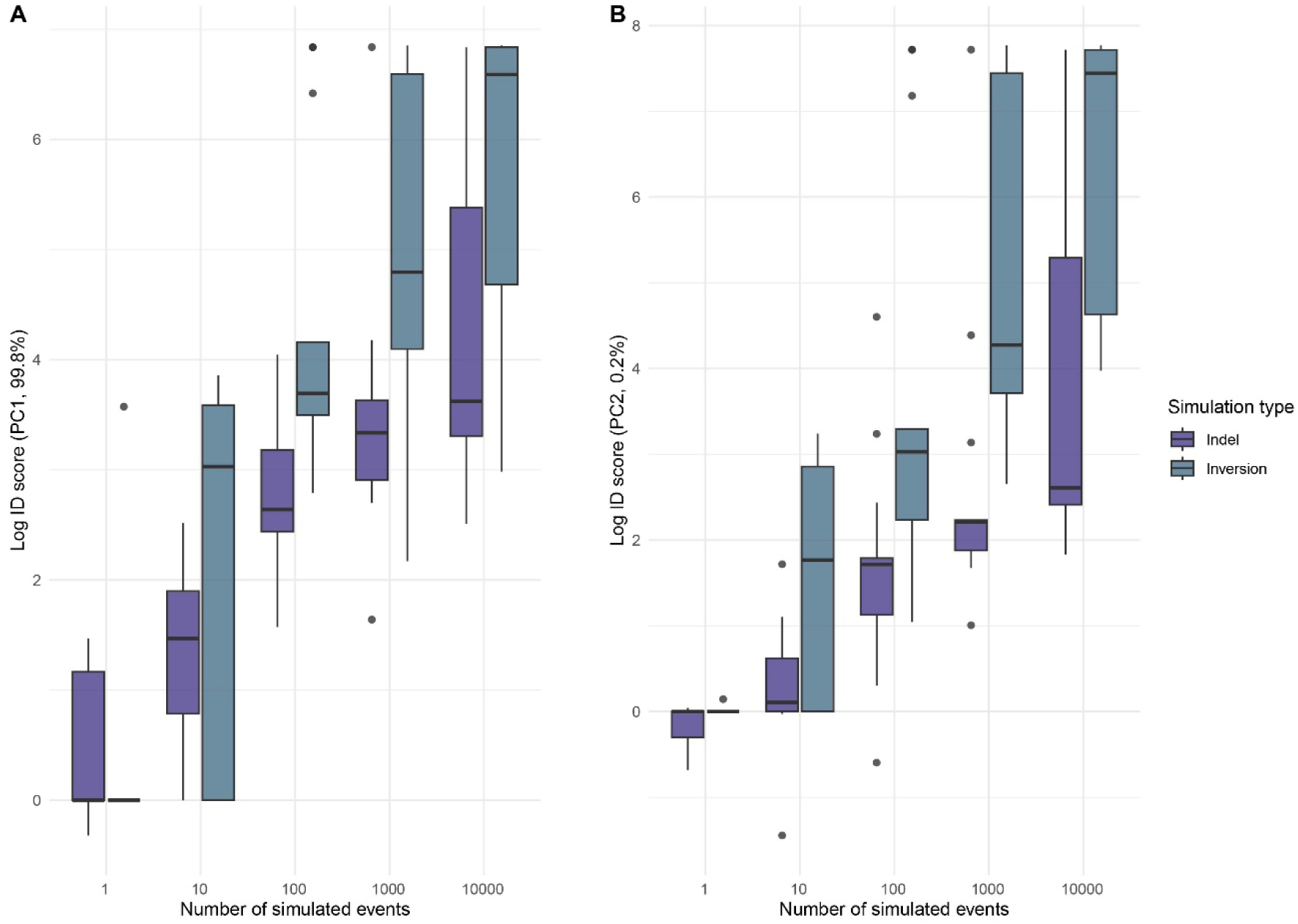
Boxplots comparing increase in disparity (ID) between indel and inversion simulations for principal component (PC)1 (A) and PC2 (B), respectively. Note that inversions cause significantly greater ID than indels.

**Figure S11.**
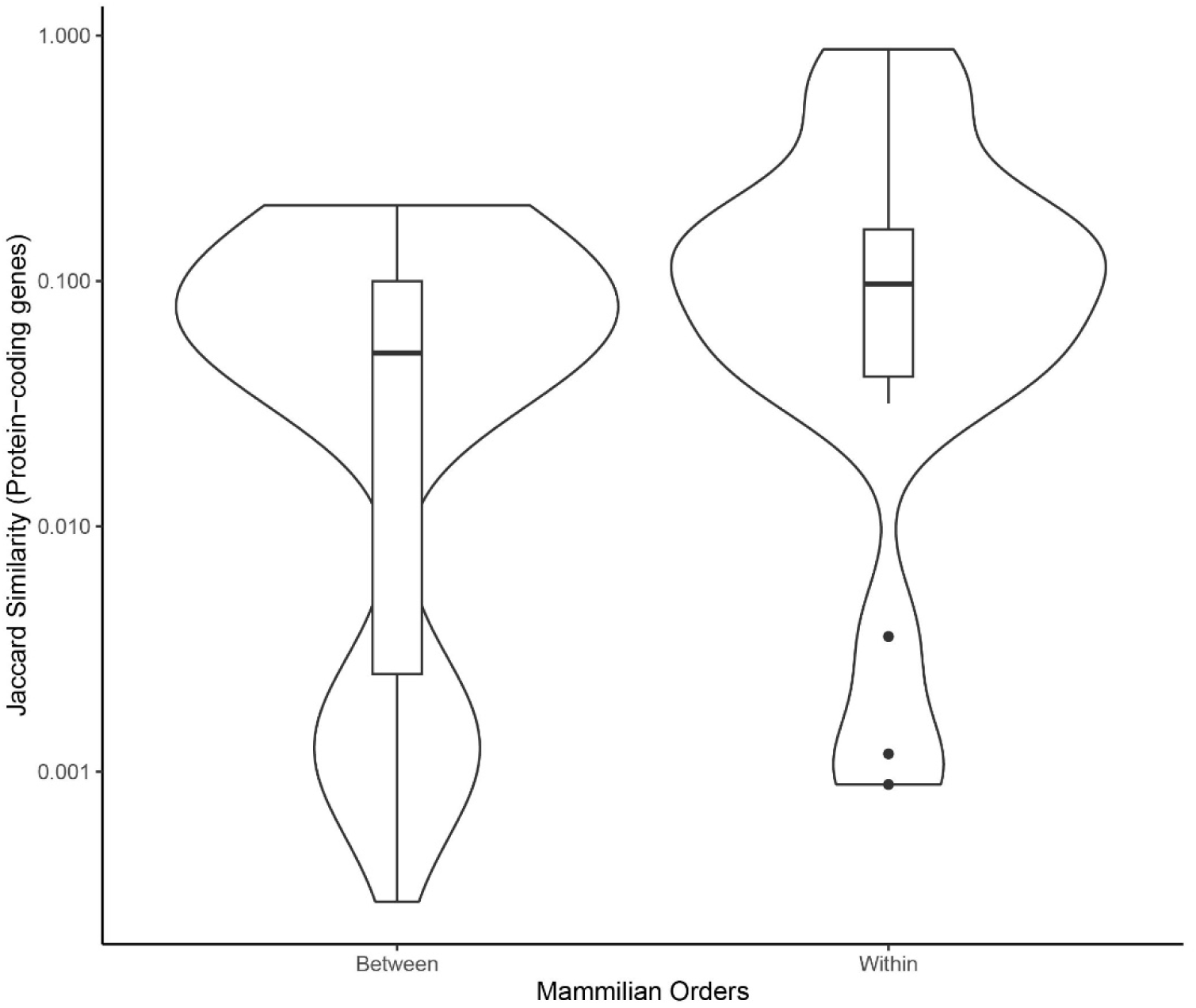
Jaccard similarity coefficients for within and between protein-coding gene content in mammalian orders used in the study.

**Figure S12.**
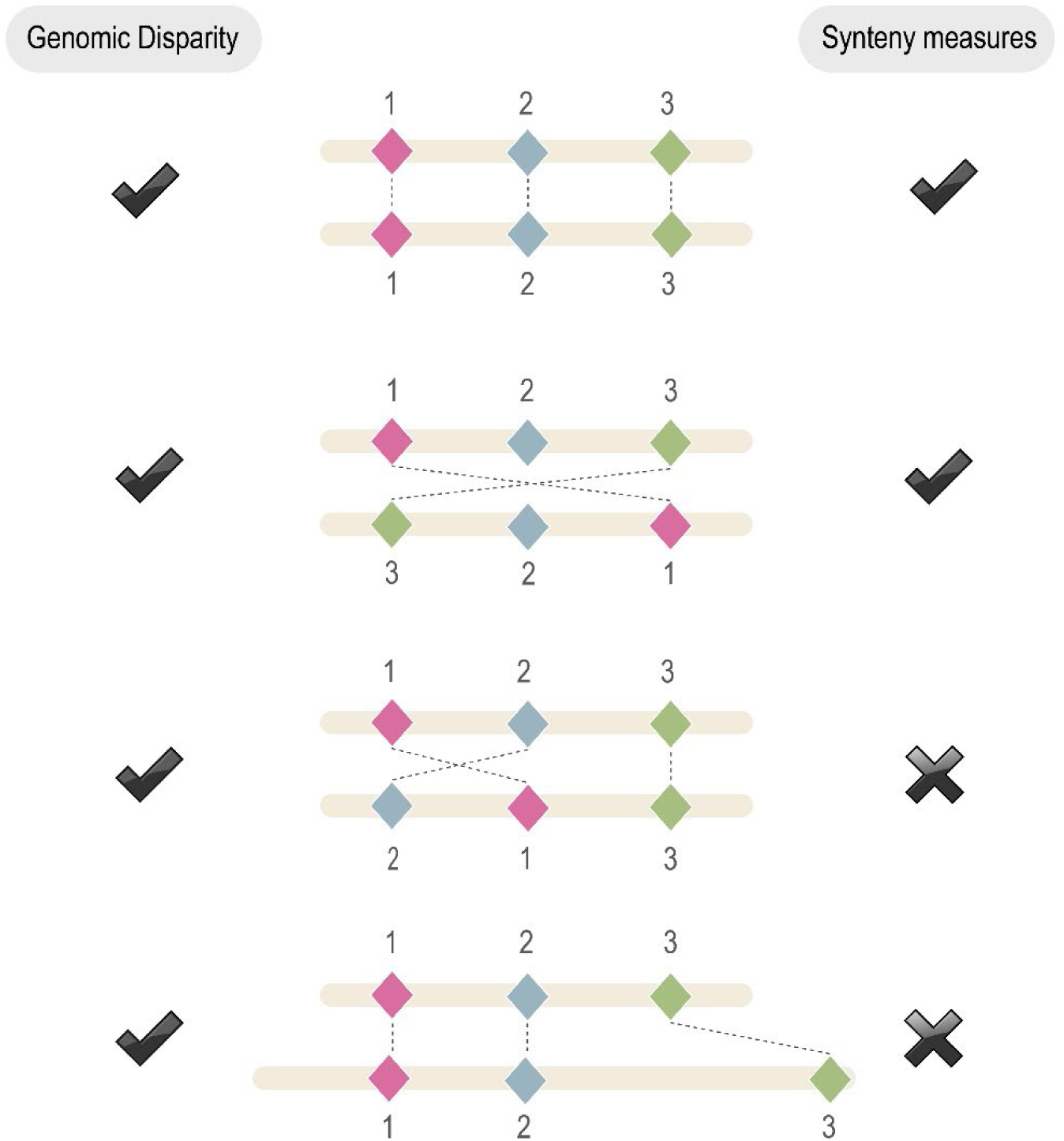
Schematic diagram illustrating sensitivities of disparity analysis and synteny analysis to different landmark scenarios. Scenarios are described from top to bottom of the figure. First, both analytical approaches will report highly conserved landmark/ortholog order, indicating when two species have highly conserved genomic architecture. Second, both approaches will report large inversions among landmark positions. Third, depending on parameter settings syntenic analysis may ignore small landmark rearrangements whereas disparity analysis captures this variation. Finally, synteny analysis ignores spatial variation in landmark placement and ‘synteny’ can be broken if chromosomal gaps are large enough whereas disparity analysis captures this variation in landmark placement.

## Appendix S1.

List of 53 BUSCO genes identified as landmarks for the human X-linked study of amniote vertebrates. eda, dlg3, snx12, gjb1, zmym3, nono, shroom4, amot, lrch2, pls3, plp1, gla, btk, drp2, sytl4, srpx2, tspan6, dach2, apool, rps6ka6, lpar4, magt1, fgf16, zdhhc15, uprt, nsdhl, vma21, cd99l2, mtmr1, mtm1, aff2, fmr1, slitrk4, atp11c, cd40lg, vgll1, htatsf1, gpc3, rap2c, frmd7, bcorl1, utp14a, smarca1, stag2, cul4b, lamp2, tmem255a

