## Supplementary Figures for "Quantifying structural variants in chromosomes using landmark-based disparity"

**This file includes:**

Figures S1 to S12

Appendix S1

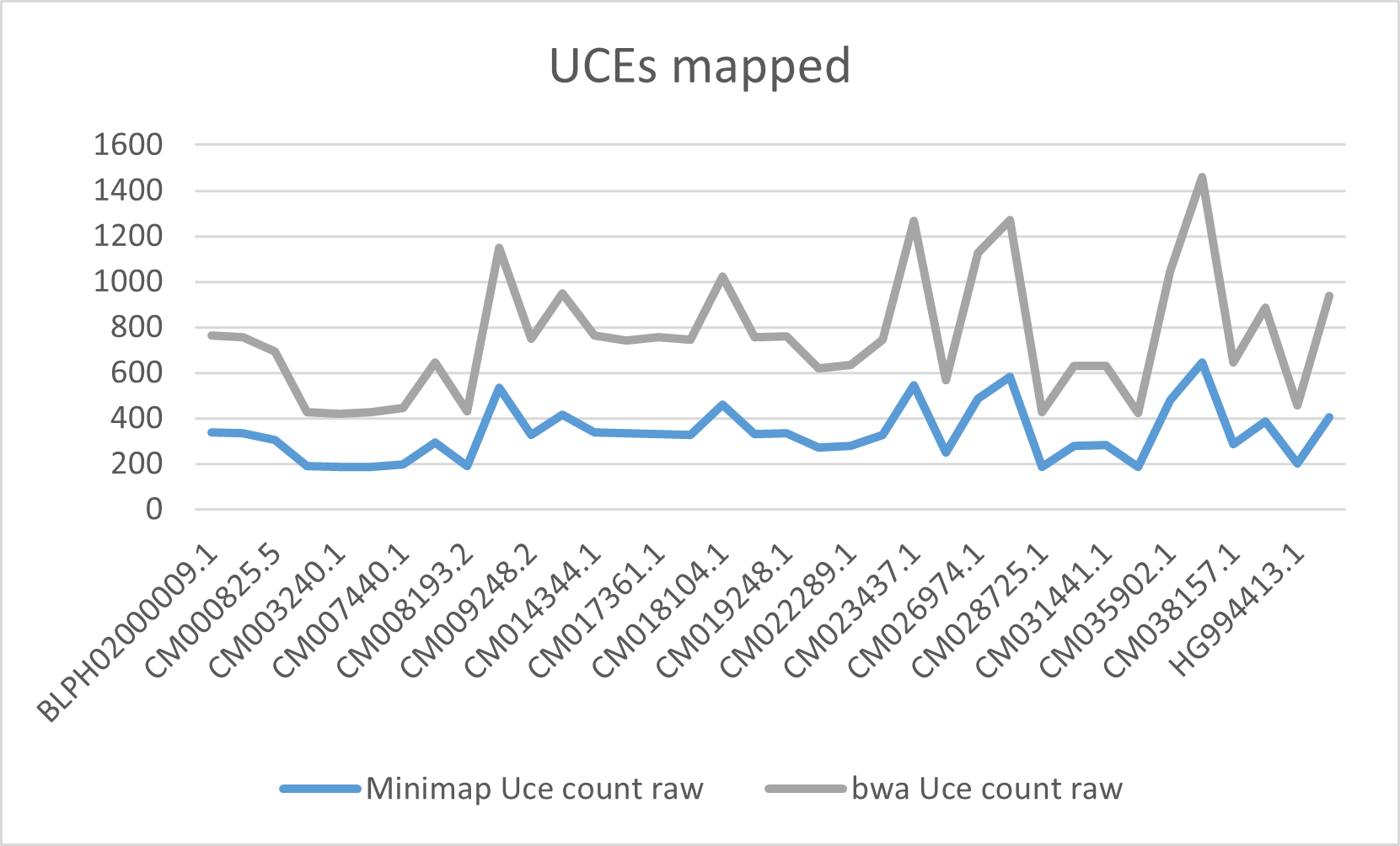

Figure S1. Number of ultraconserved elements (UCEs) mapped utilizing two different mapping software: minimap2 (blue) and bwa-mem (grey). Demonstrated using a subset of chromosome assemblies picked from the ‘chromosome set 2’ of the placental mammals UCE dataset discussed in the main text.

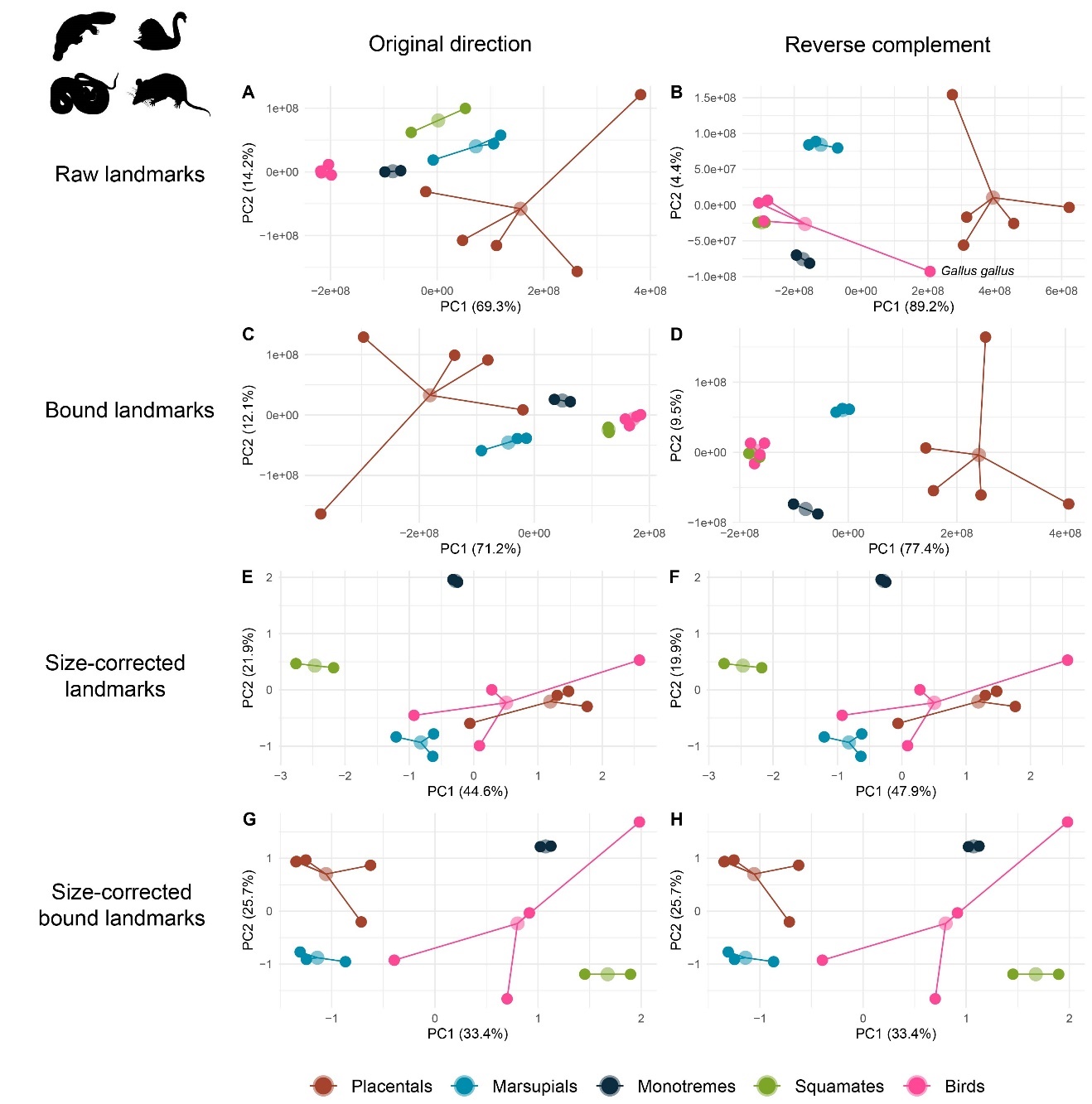

Figure S2. Principal component (PC) sensitivity analyses of 53 X-linked BUSCO landmarks in amniote vertebrates taken from Lovell et al. (2022). The first row are PCAs conducted using raw, absolute positions for the original direction (A) and reverse complement (B), respectively. The second row are PCAs conducted using bounded, absolute positions for the original direction (C) and the reverse complement (D). The third row are PCAs conducted using size-correction with chromosome size for the original direction (E) and the reverse complement (F). The fourth row of plots are PCAs conducted using size-correction using bounded chromosome sizes and positions for the original direction (G) and reverse complement (H). The chicken (*Gallus gallus*) is highlighted in plot B to illustrate an analytical artifact associated with clustered landmarks on large chromosomes. Organism silhouettes downloaded from <https://www.phylopic.org/>.

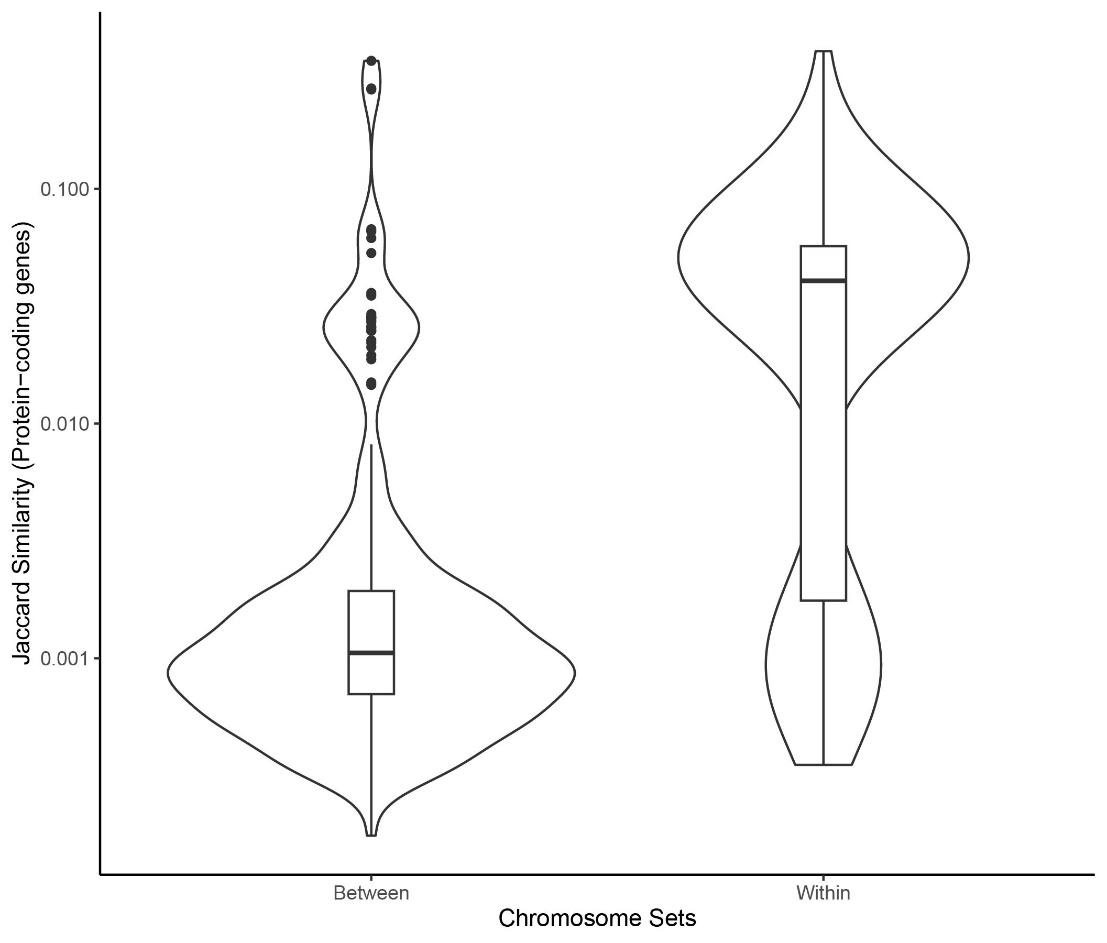

Figure S3. Jaccard similarity coefficients for within and between protein-coding gene content in the two focal chromosome sets used in the study.

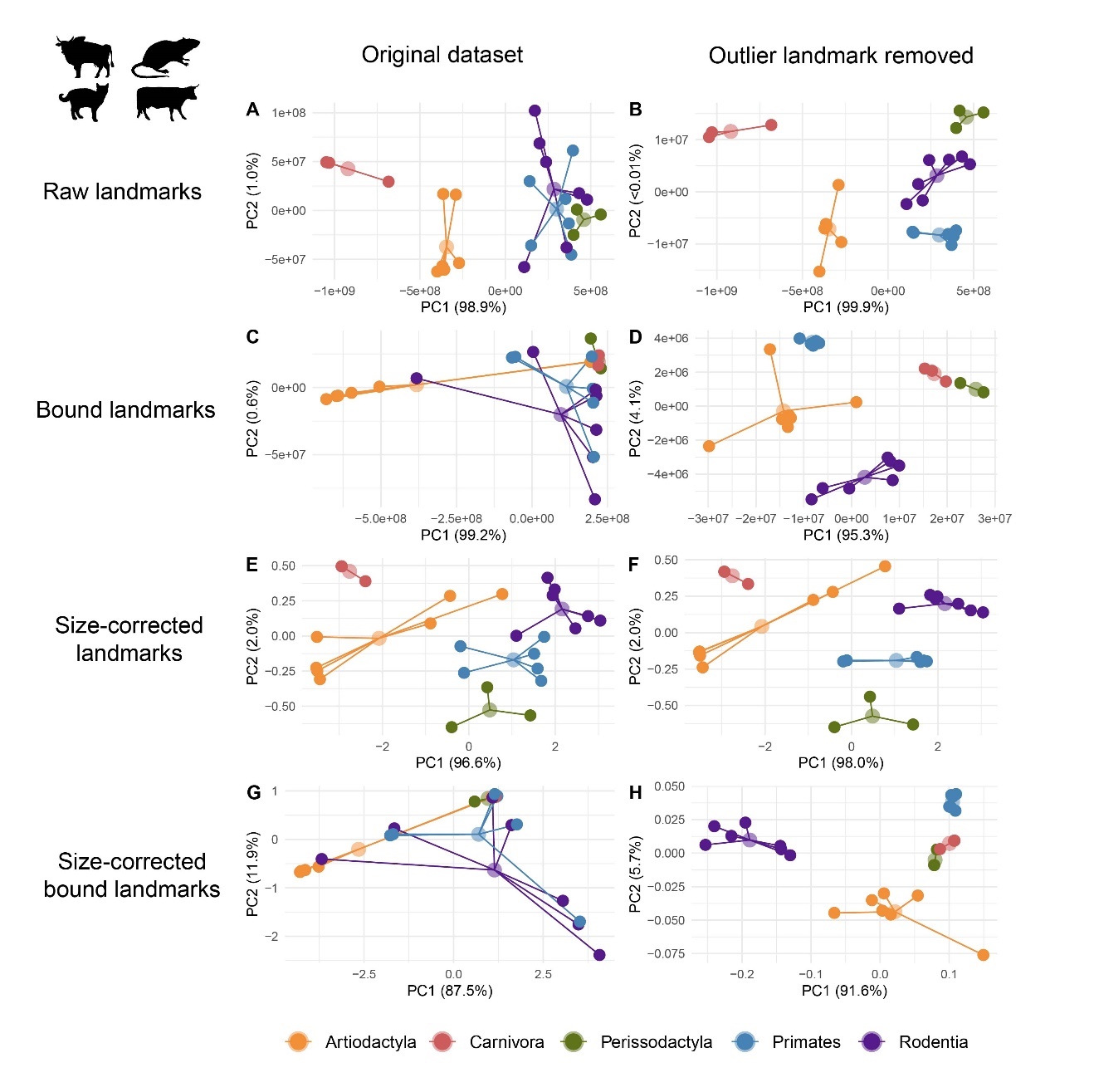

**Figure S4.** Principal component analyses (PCAs) of ultraconserved element (UCE) landmarks in placental mammals for chromosome set 1 before and after the removal of an outlier landmark (UCE 6948). The first row of plots are PCAs conducted using raw, absolute positions for the original direction (A) and reverse complement (B), respectively. The second row of plots are PCAs conducted using bounded, absolute positions for the original direction (C) and the reverse complement (D). The third row of plots are PCAs conducted using size-correction with chromosome size for the original direction (E) and the reverse complement (F). The fourth row of plots are PCAs conducted using size-correction using bounded chromosome sizes and positions for the original direction (G) and reverse complement (H). The first column of plots are PCAs conducted with all 186 UCE landmarks and the second column of plots are PCAs based on 185 UCE landmarks (outlier removed).

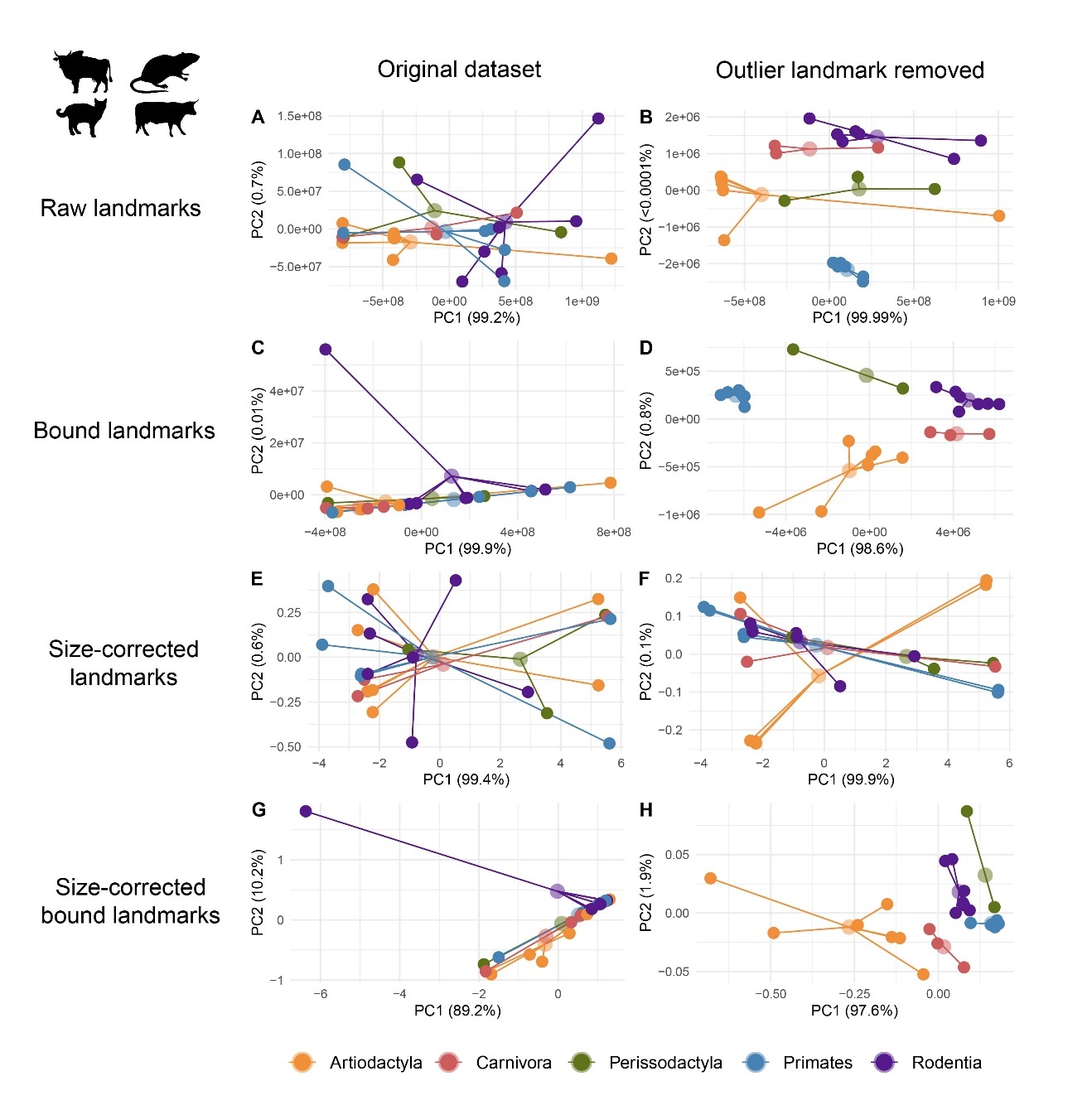

**Figure S5.** Principal component analyses (PCAs) of ultraconserved element (UCE) landmarks in placental mammals for chromosome set 2 before and after the removal of an outlier landmark (UCE 6948). The first row of plots are PCAs conducted using raw, absolute positions for the original direction (A) and reverse complement (B), respectively. The second row of plots are PCAs conducted using bounded, absolute positions for the original direction (C) and the reverse complement (D). The third row of plots are PCAs conducted using size-correction with chromosome size for the original direction (E) and the reverse complement (F). The fourth row of plots are PCAs conducted using size-correction using bounded chromosome sizes and positions for the original direction (G) and reverse complement (H). The first column of plots are PCAs conducted with all 82 UCE landmarks and the second column of plots are PCAs based on 81 UCE landmarks (outlier removed).

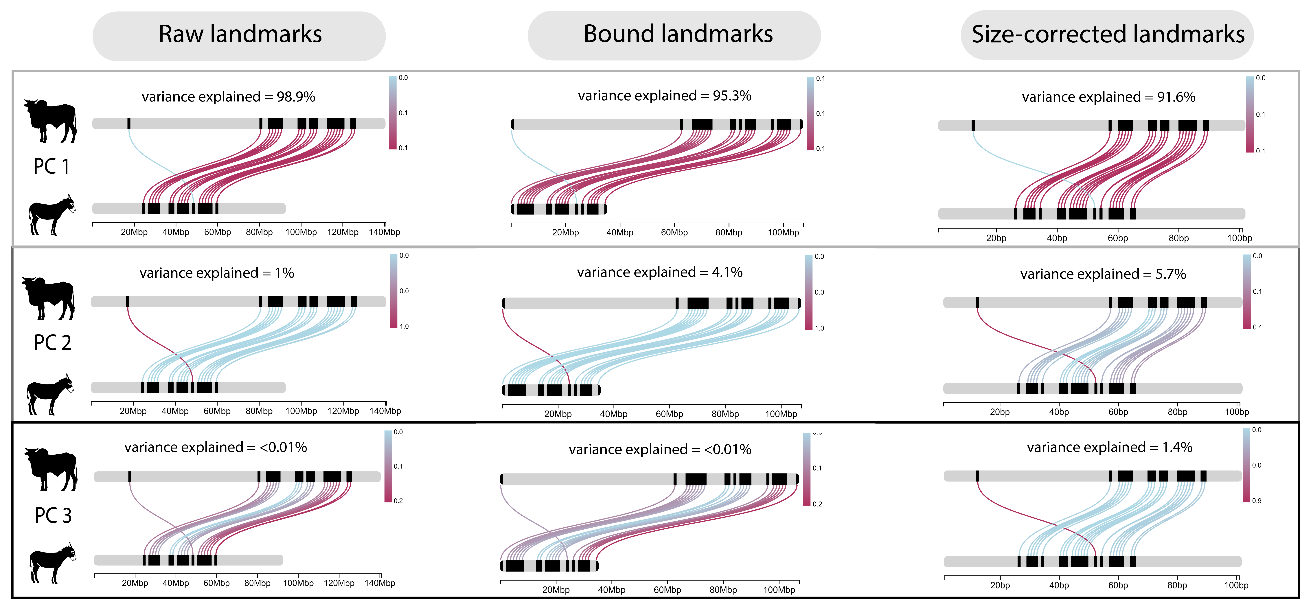

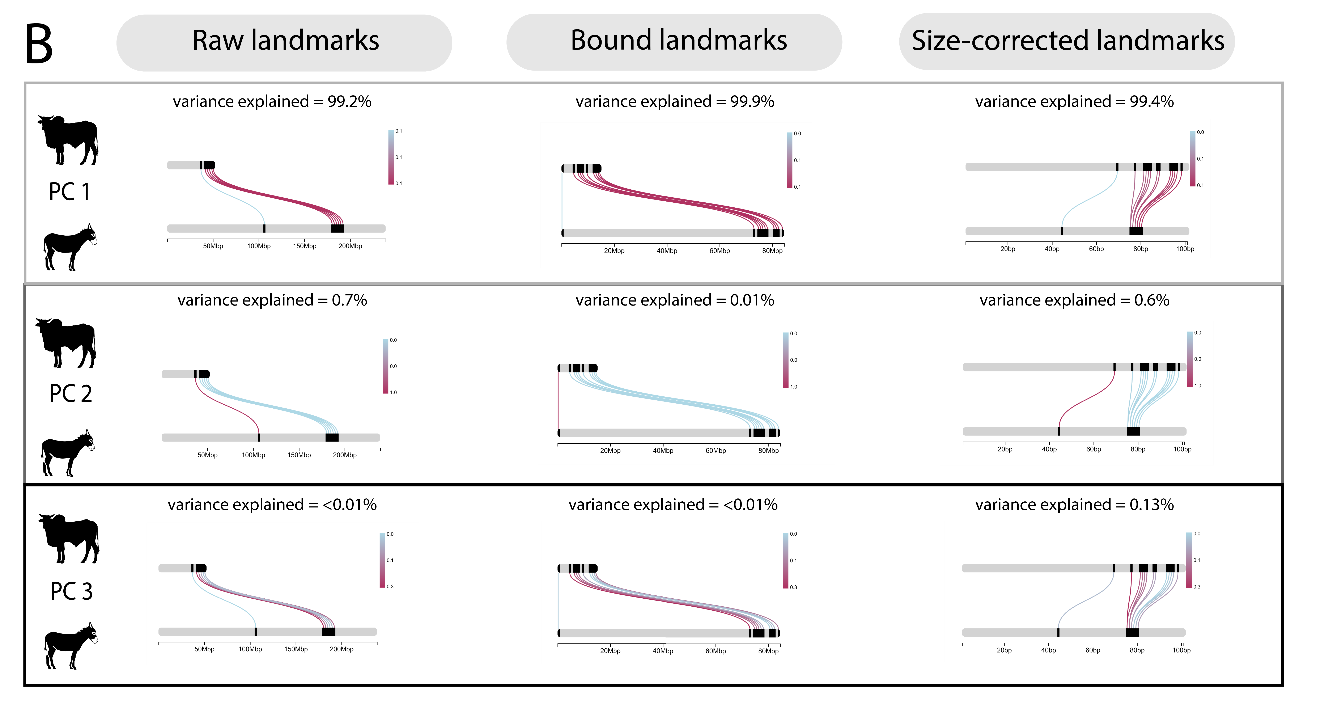

**Figure S6.** ChromoMap plots visualizing principal component analysis (PCA) loadings of landmarks on PC1, PC2, and PC3 in the raw landmark PCA, bounded landmark PCA, and size corrected landmark PCA of chromosome set 1 (A) and set 2 (B) from the placental mammal visualized using links between *Bos indicus* and *Equus asinus*. Respective variances explained by each principal component are listed in the plot. A single landmark (UCE 6948) was found to be uniquely weighed on the various PCs identifying it as an outlier landmark.

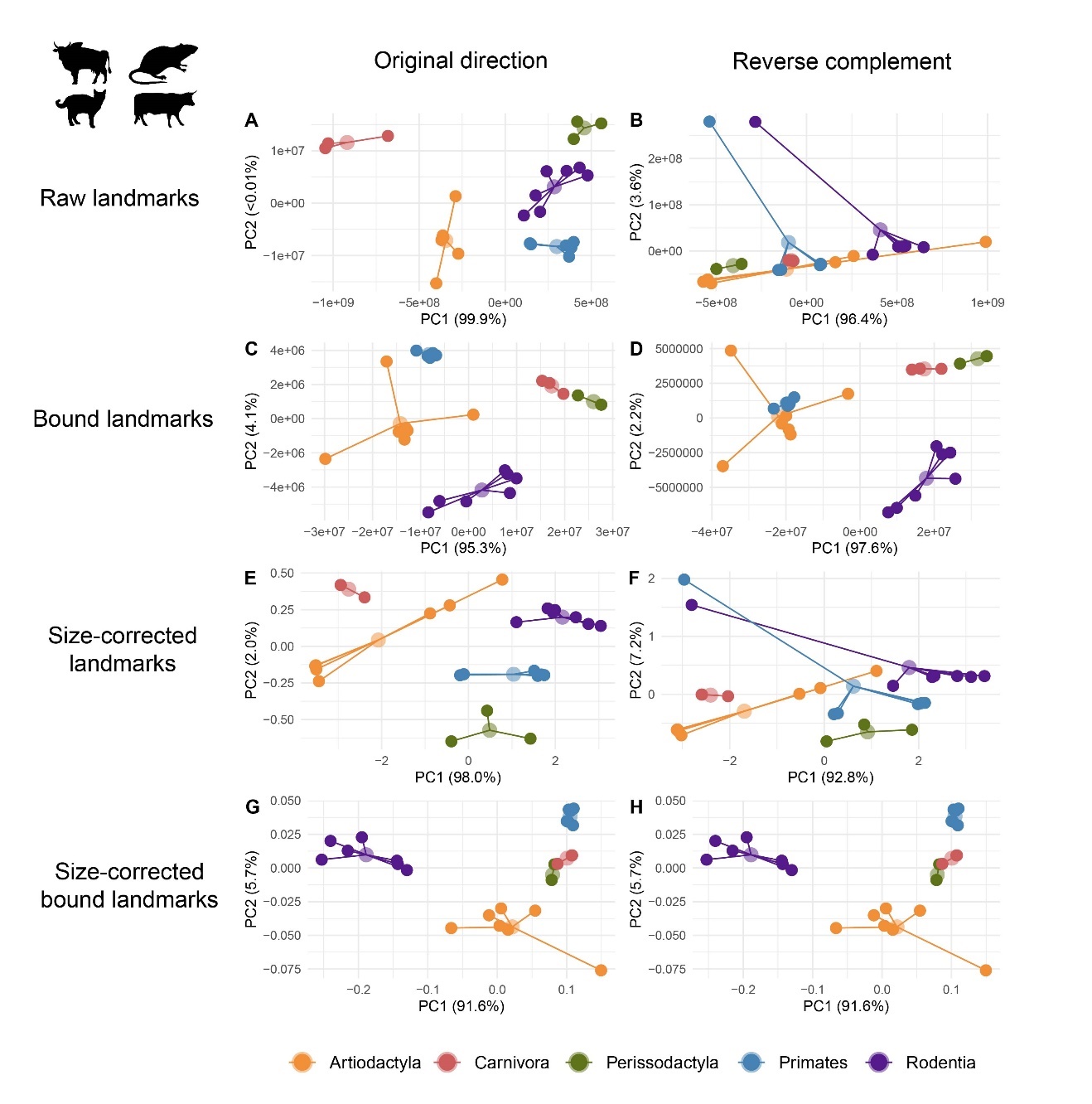

**Figure S7.** Principal component (PC) sensitivity analyses of outlier removed UCE landmarks in placental mammals chromosome set 1. The first row are PCAs conducted using raw, absolute positions for the original direction (A) and reverse complement (B), respectively. The second row are PCAs conducted using bounded, absolute positions for the original direction (C) and the reverse complement (D). The third row are PCAs conducted using size-correction with chromosome size for the original direction (E) and the reverse complement (F). The fourth row are PCAs conducted using size-correction using bounded chromosome sizes and positions for the original direction (G) and reverse complement (H).

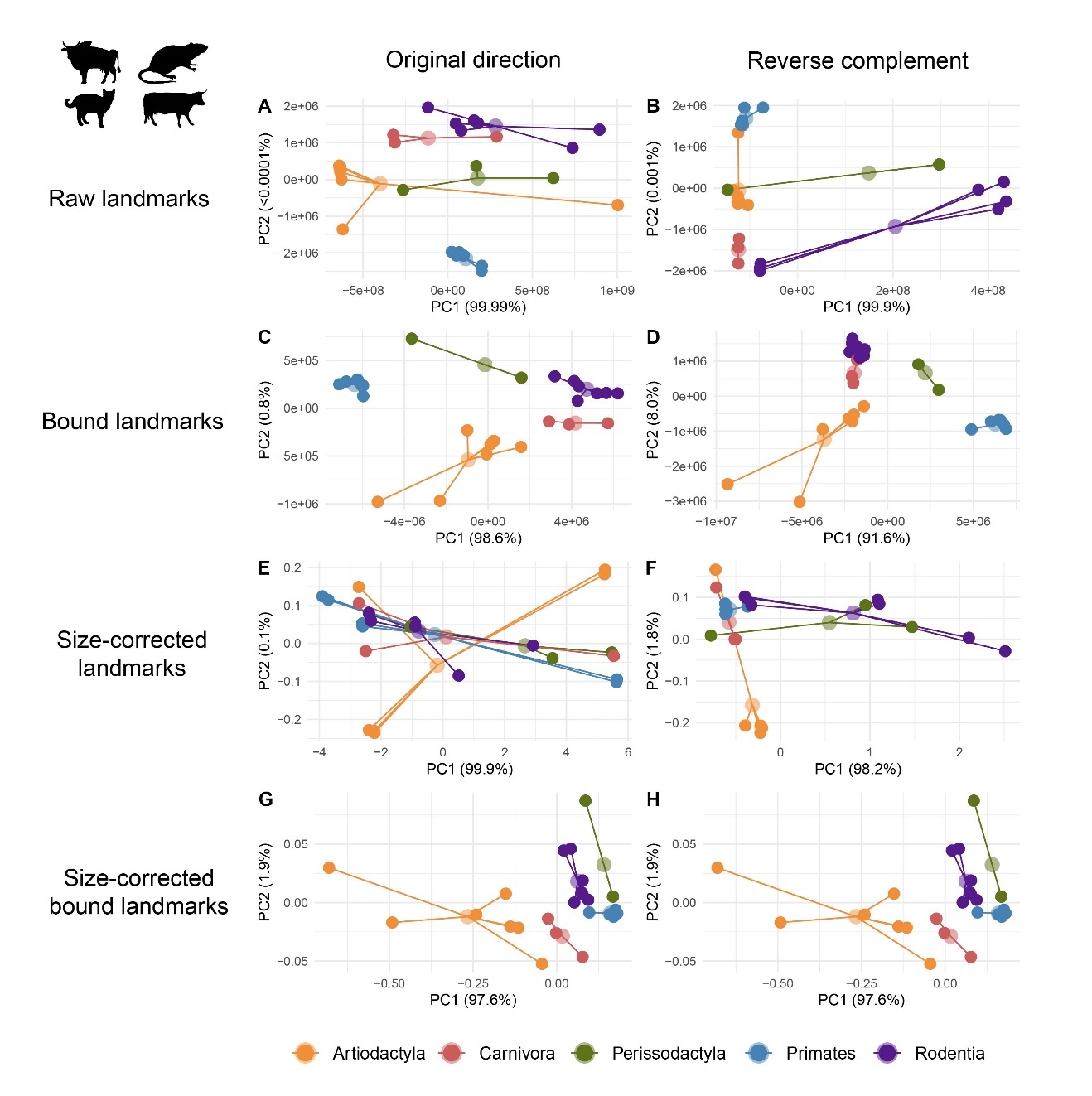

**Figure S8.** Principal component (PC) sensitivity analyses of outlier removed UCE landmarks in placental mammals chromosome set 2. The first row are PCAs conducted using raw, absolute positions for the original direction (A) and reverse complement (B), respectively. The second row are PCAs conducted using bounded, absolute positions for the original direction (C) and the reverse complement (D). The third row are PCAs conducted using size-correction with chromosome size for the original direction (E) and the reverse complement (F). The fourth row are PCAs conducted using size-correction using bounded chromosome sizes and positions for the original direction (G) and reverse complement (H).

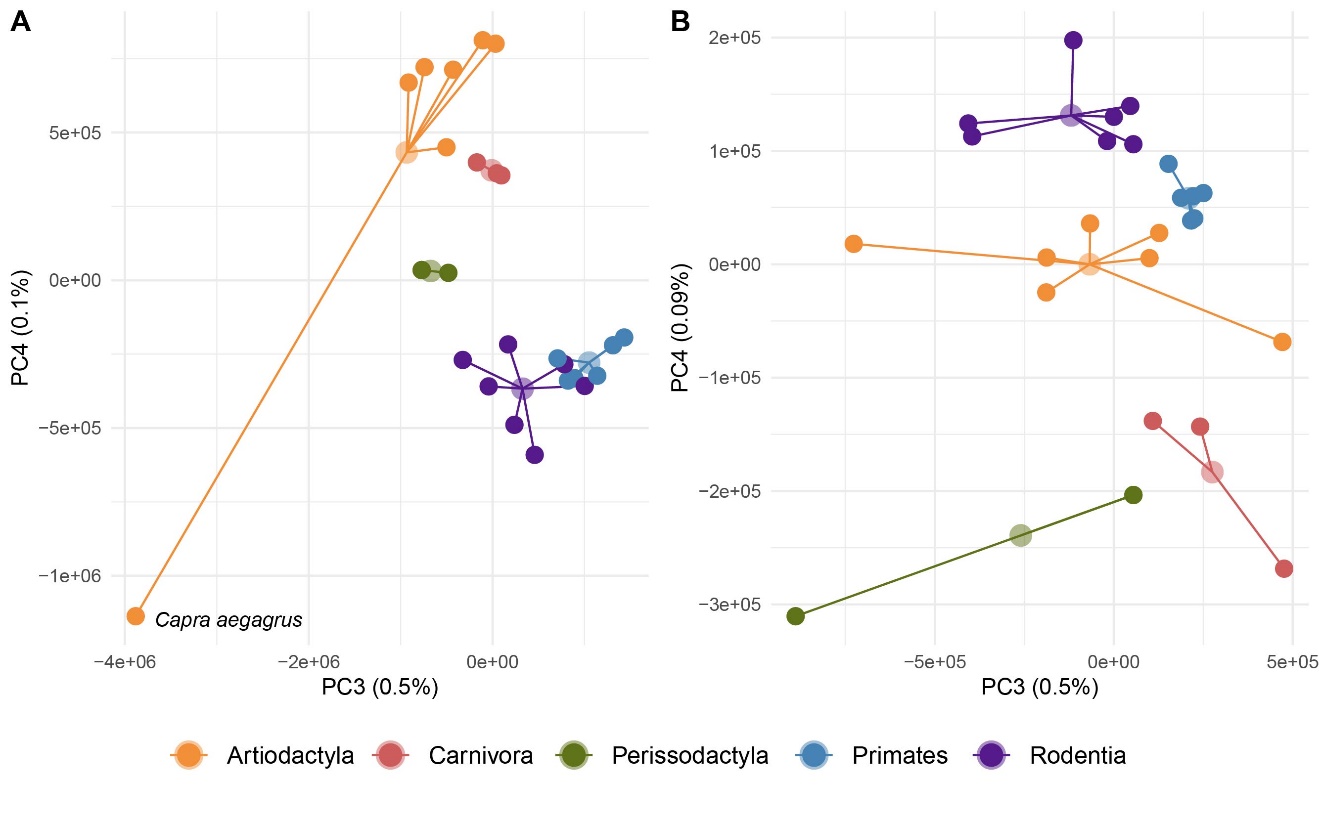

**Figure S9**. Ordination results from principal component (PC) analysis of UCE landmark positions in two sets of chromosomes from 26 species of placental mammals using bounded raw positions after the removal of an outlier landmark (see text). Representatives for five orders (Artiodactyla, Carnivora, Perissodactyla, Primates and Rodents) were included (Table1). The depicted structural variation is inferred from PCs 3 and 4 of 185 (chromosome set 1, A) and 81 (chromosome set 2, B) landmarks, respectively. In both datasets, the first two PCs explained > 99% of the variation (see **Figure 5**, main text). While explaining < 1% of the overall variation, PCs 3 and 4 mostly separated mammalian orders in geno-metric space. *Carpa aegragus* was a notable outlier in chromosome set 1.

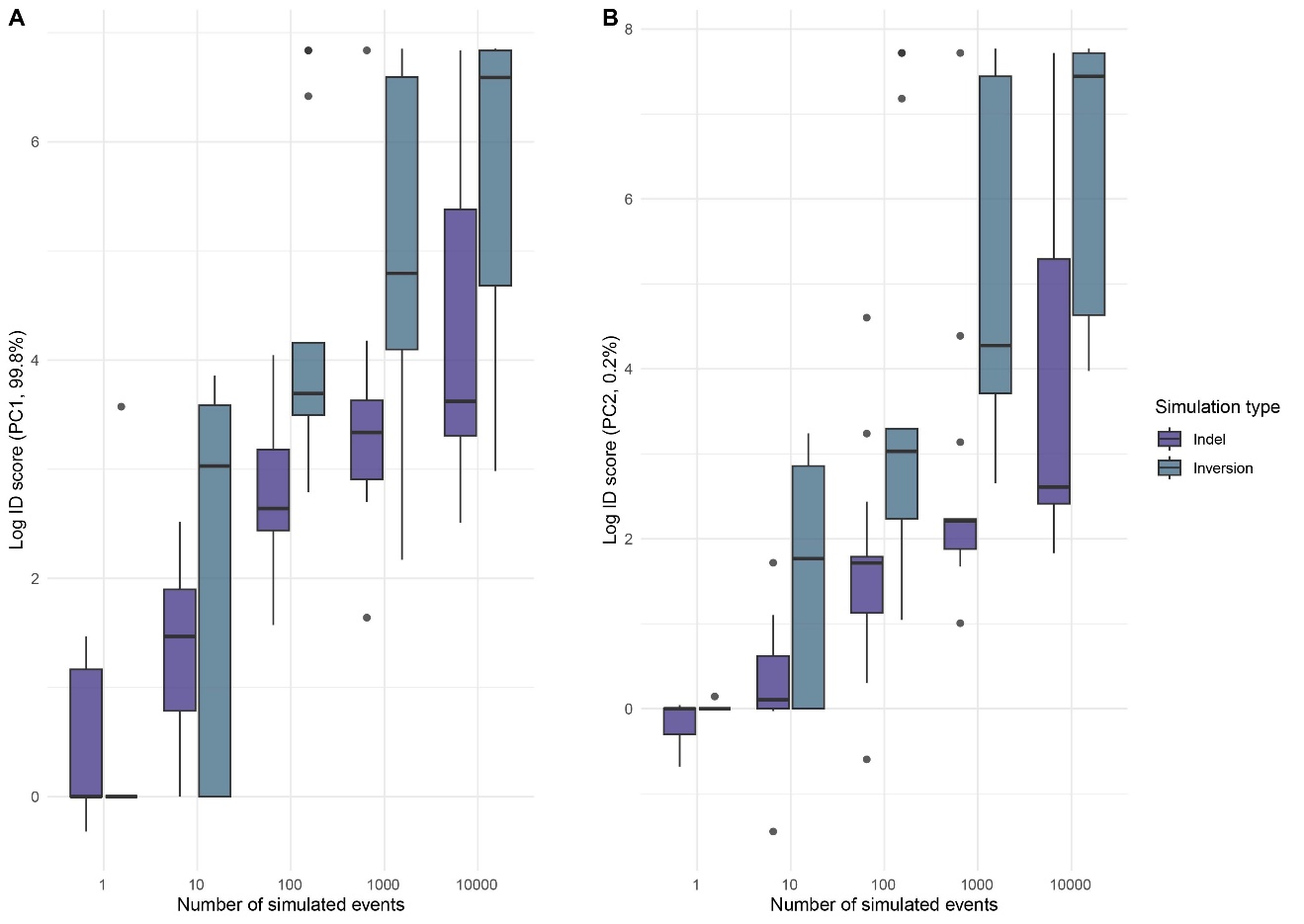

**Figure S10.** Boxplots comparing increase in disparity (ID) between indel and inversion simulations for principal component (PC)1 (A) and PC2 (B), respectively. Note that inversions cause significantly greater ID than indels.

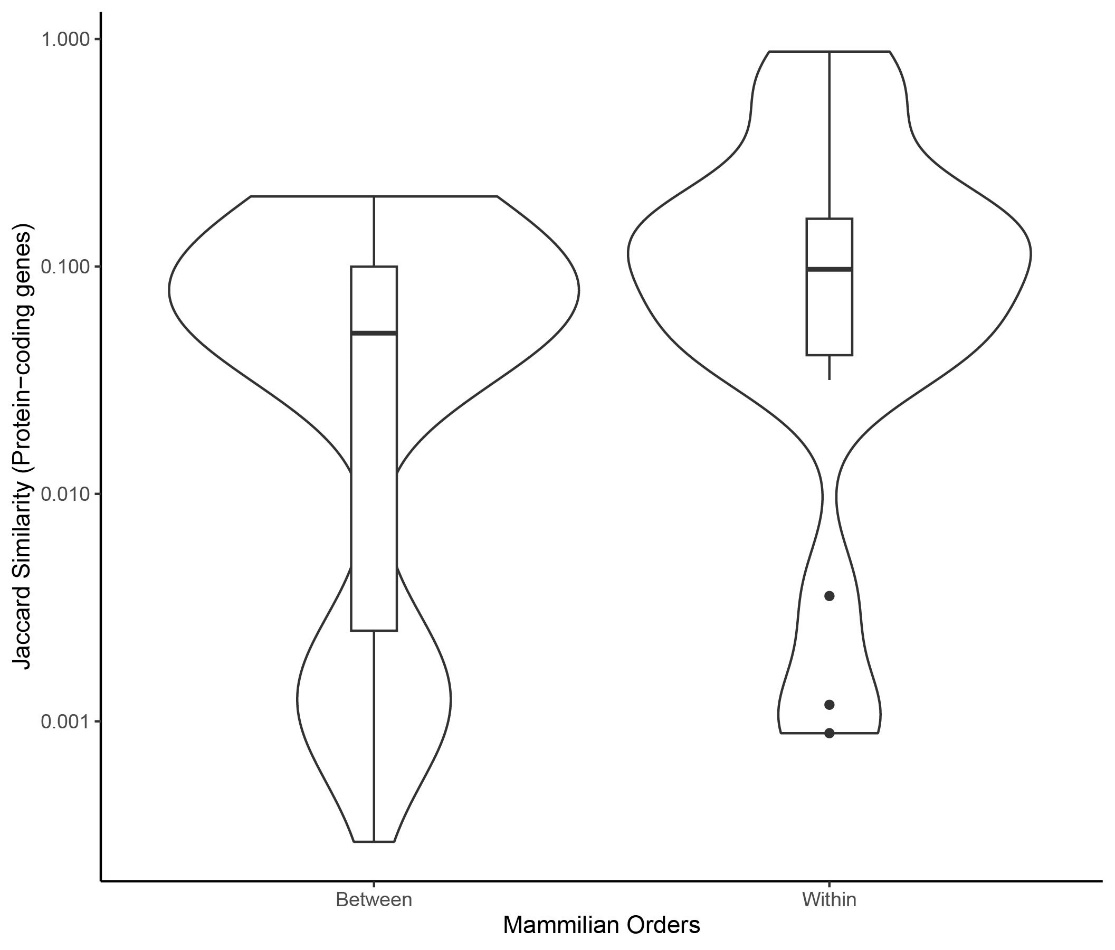

**Figure S11.** Jaccard similarity coefficients for within and between protein-coding gene content in mammalian orders used in the study.

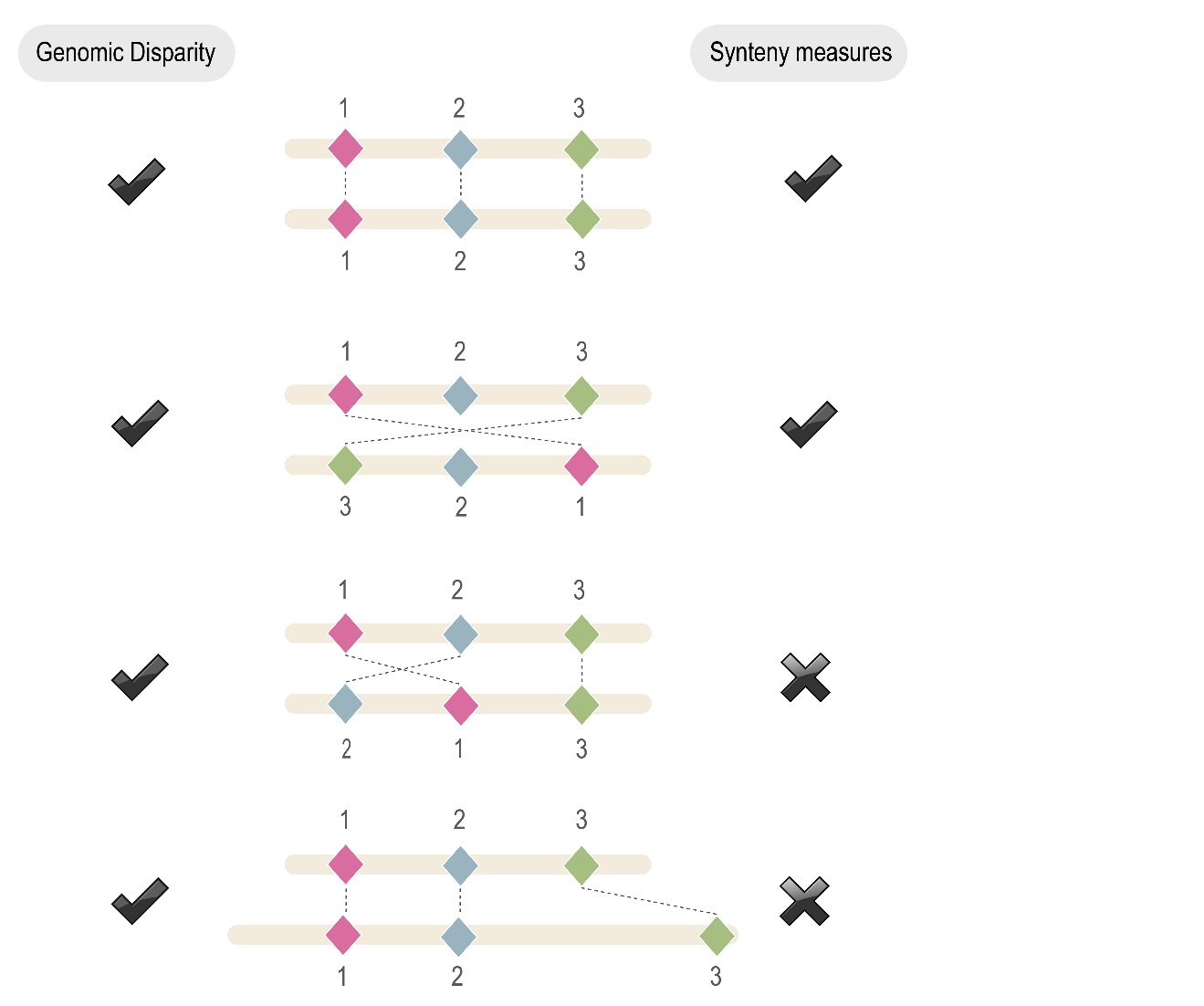

**Figure S12.** Schematic diagram illustrating sensitivities of disparity analysis and synteny analysis to different landmark scenarios. Scenarios are described from top to bottom of the figure. First, both analytical approaches will report highly conserved landmark/ortholog order, indicating when two species have highly conserved genomic architecture. Second, both approaches will report large inversions among landmark positions. Third, depending on parameter settings syntenic analysis may ignore small landmark rearrangements whereas disparity analysis captures this variation. Finally, synteny analysis ignores spatial variation in landmark placement and ‘synteny’ can be broken if chromosomal gaps are large enough whereas disparity analysis captures this variation in landmark placement.

**Appendix S1.** List of 53 BUSCO genes identified as landmarks for the human X-linked study of amniote vertebrates.

eda, dlg3, snx12, gjb1, zmym3, nono, shroom4, amot, lrch2, pls3, plp1, gla, btk, drp2, sytl4, srpx2, tspan6, dach2, apool, rps6ka6, lpar4, magt1, fgf16, zdhhc15, uprt, nsdhl, vma21, cd99l2, mtmr1, mtm1, aff2, fmr1, slitrk4, atp11c, cd40lg, vgll1, htatsf1, gpc3, rap2c, frmd7, bcorl1, utp14a, smarca1, stag2, cul4b, lamp2, tmem255a, zbtb33, upf3b, pgrmc1, fam199x, il1rapl2, arhgef9
